# A universal law of thermodynamics-kinetics coupling shapes enzyme allocation and glycolytic efficiency across species

**DOI:** 10.1101/2025.09.29.679393

**Authors:** Kai Sun, Weiyan Zheng, Wenchao Fan, Chuyun Ding, Dan Huang, Ziwei Dai

## Abstract

Efficient use of limited cellular resources is fundamental to metabolism. Although flux optimization is widely recognized as a central objective of metabolic networks, how flux efficiency influences the allocation of the metabolic proteome remains unclear and lacks direct validation. Here, we derive a simple analytical relationship linking the equilibrium constant and the catalytic-abundance quotient of reactions within a pathway that defines the condition for maximal efficiency. We refer to this principle as the law of thermodynamics-kinetics coupling. By integrating reaction thermodynamics, enzyme kinetics, transcriptomic, and proteomic data, we show that glycolytic enzyme allocation consistently obeys this law across evolutionarily distant species. Moreover, the drive to optimize glycolytic efficiency is strengthened under oncogenic signaling and limited cellular budget for glycolytic enzymes. These findings establish a universal principle governing enzyme allocation in metabolic pathways and reveal key determinants of efficiency optimality in glycolysis.

**Significance statement:** Current models of metabolic optimality either rely on coarse-grained proteome allocation or constraint-based simulations, largely neglecting the fundamental constraints imposed by enzyme kinetics and thermodynamics on pathway efficiency. Here, we derive and validate a universal law of thermodynamics-kinetics coupling that robustly governs the allocation of glycolytic enzymes across species under optimal efficiency. This study has three major advances: it reveals universal economic principles of metabolism with simple, generalized mathematical form, uncovers determinants underlying the heterogeneity of metabolic efficiency optimization, and provides a practical blueprint for efficiency optimization of engineered pathways.

## Introduction

From autocatalytic chemical reaction systems in the prebiotic world to complex, compartmentalized metabolic networks of modern cells, the evolution of metabolism has lasted for billions of years. The optimization of fitness during evolution of metabolism covers multiple facets, including its ability to sustain robust metabolic fluxes, minimize investment of resources, and maintain functions upon environmental perturbations. As one of the earliest biological processes to be mathematically modeled, these metabolic objectives can be formalized to predict metabolic states that maximize fitness^1^.

Efficient resource utilization is a central goal of metabolism. Among essential resources of metabolism, the cellular proteome is a major investment, as biosynthesis of proteins is among the most energy-consuming cellular processes, contributing to roughly 20% of the basal metabolic rate^2^. Consequently, the proteome is under strong pressure of efficiency optimization. Evidence for this theory spans multiple scales: amino acids with low synthetic costs are preferred in the proteome of unicellular organisms such as bacteria^3, 4^ and yeast^5^ as well as in cancer cells^6^. Dietary amino acid intake matching the proteomic amino acid composition has beneficial health outcomes in multicellular organisms^7^, probably by saving the cost of amino acid biosynthesis. The goal of minimizing proteome cost is implicitly encoded in the current codon system which serves as the very fundamental logic of almost all existing organisms to minimize the potential increase in amino acid biosynthetic cost under mutations^8^. At the whole proteome level, coarse-grained mathematical modeling and quantitative biology experiments have shown that its allocation into major functional groups such as ribosome and metabolism for maximizing proliferation can precisely explain the emergence of growth scaling law, overflow metabolism, and many other phenomena in several bacteria species^9-12^.

A key question about such economic principle is how it governs the operation of individual metabolic pathways. Proteomics data across bacteria species showed that relative abundance of glycolytic enzymes are conserved across species and related to reaction thermodynamics, implying existence of such optimality principles^13^. This can be mathematically framed as maximizing pathway flux under limited enzyme abundance - a constrained optimization problem solvable either analytically or numerically if reaction kinetics and thermodynamics are known. While theoretical works on optimal proteome allocation often adopt a coarse-grained strategy focusing on major functional classes, our understanding about optimal enzyme allocation within individual pathways is still limited. Early theoretical work of metabolic control analysis (MCA) derived general conditions for optimal efficiency^14-16^. For instance, upon optimal pathway efficiency, the flux control coefficients (FCCs), which quantify how the pathway flux responds to perturbations of single enzymes, are proportional to the enzyme abundances^14^. However, a major challenge is translating these theoretical principles into testable predictions against modern omics data, because the theory of MCA was developed in 1970s, long before high throughput sequencing-based omics technologies were available. A numerical method for solving the optimal enzyme allocation^17^ and rate-law-dependent analytical solutions have been derived later, but for realistic Michaelis-Menten kinetics, analytical solution is unavailable as the optimization problem becomes highly nonlinear^18^. Moreover, these theoretical models ignore multi-substrate reactions and isozyme-catalyzed reactions, hence their predictions are difficult to validate robustly due to the noise in experimental measurements and complexity of in-vivo metabolic systems. Therefore, a universal, simple, and experimentally testable quantitative law connecting enzyme abundance and reaction characteristics at optimal efficiency is lacking.

In this study, we establish a universal law of thermodynamics-kinetics coupling for linear pathways at optimal efficiency. This law provides a simple, explicit relationship between reaction equilibrium constants, catalytic efficiencies, and enzyme abundances testable with real-world reaction kinetics, thermodynamics, and omics data. Using glycolysis as a model pathway, we validate the law across bacteria, yeast, and human cells. We quantified the tendency of efficiency optimization for glycolysis and found that stronger efficiency optimization is associated with enhanced oncogenic signaling in tumors and limited total amount of glycolysis enzymes across diverse conditions. These findings reveal that a simple economic principle of efficiency optimization quantitatively governs enzyme allocation in metabolism across phylogenetically distant species.

## Results

### Allocation of human metabolic enzyme abundances is conserved

To test the hypothesis that metabolic enzyme abundance follows conserved allocation principles, we analyzed quantitative proteomics data for 378 human cancer cell lines in the CCLE project^19^ and transcriptomics data for TCGA tumors and normal tissues. We found that the allocation of glycolytic enzymes was highly conserved across cell lines with different tissues-of-origin (Figure 1a). The conservation persists when extended to genome-scale metabolic enzymes, with median pairwise Spearman’s rank correlation coefficient of metabolic enzyme abundance exceeding 0.8 at both protein and transcript levels (Fig. 1b-e). A moderate correlation (Spearman correlation around 0.5) was also observed between protein abundances in the cell lines and transcript levels in the tumors (Fig. S1), suggesting that the allocation of metabolic enzyme abundance is robust across different omics layers. These results reveal a conserved allocation of metabolic enzymes across diverse human cell and tissue types, indicating optimality principles underlying them.

**Figure 1.**
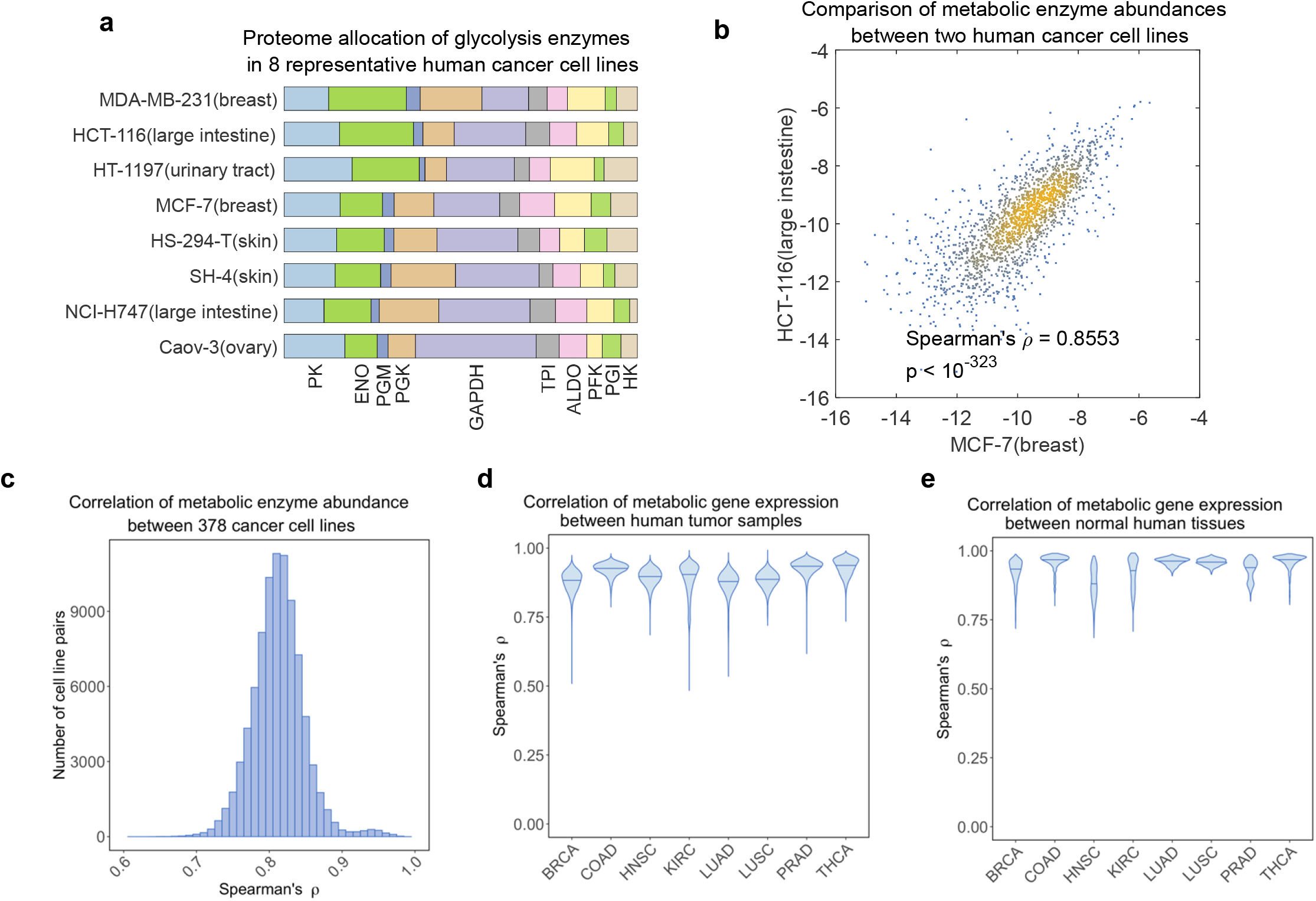
Allocation of human metabolic enzyme abundances is conserved. (a) Allocation of glycolysis enzyme abundances in 8 human cancer cell lines in the CCLE project. (b) Scatter plot comparing the abundances of metabolic enzymes between two human cancer cell lines, MCF-7 and HCT-116. (c) Distribution of Spearman’s rank correlation coefficient of metabolic enzyme abundance between 378 different human cancer cell lines in the CCLE project. (d) Distributions of Spearman’s rank correlation coefficient of metabolic gene expression between normal human tissue samples in the TCGA project. (e) Distributions of Spearman’s rank correlation coefficient of metabolic gene expression between human tumor samples in the TCGA project.

### Law of kinetics-thermodynamics coupling underlying optimal efficiency of metabolic pathways

We hypothesize that the conserved allocation of metabolic enzymes is a consequence of efficiency maximization. Efficiency of a metabolic pathway means the ability of it to achieve as high steady state flux as possible with limited total amount of metabolic enzymes. To simplify mathematical derivation, we start from the simplest linear metabolic pathway consisting of a chain of subsequent unimolecular reactions. We assume first-order reaction kinetics (Fig. 2a). At steady state, when the concentrations of boundary metabolites *S*_*in*_ and *S*_*out*_ are fixed, the pathway flux *J* is a function of enzyme abundances [*E*_*i*_], catalytic efficiencies *a*_*i*_, and equilibrium constants *K*_*i*_. We define the flux efficiency of the pathway as the ratio of the flux *J* to the total enzyme abundance 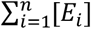:

**Figure 2.**
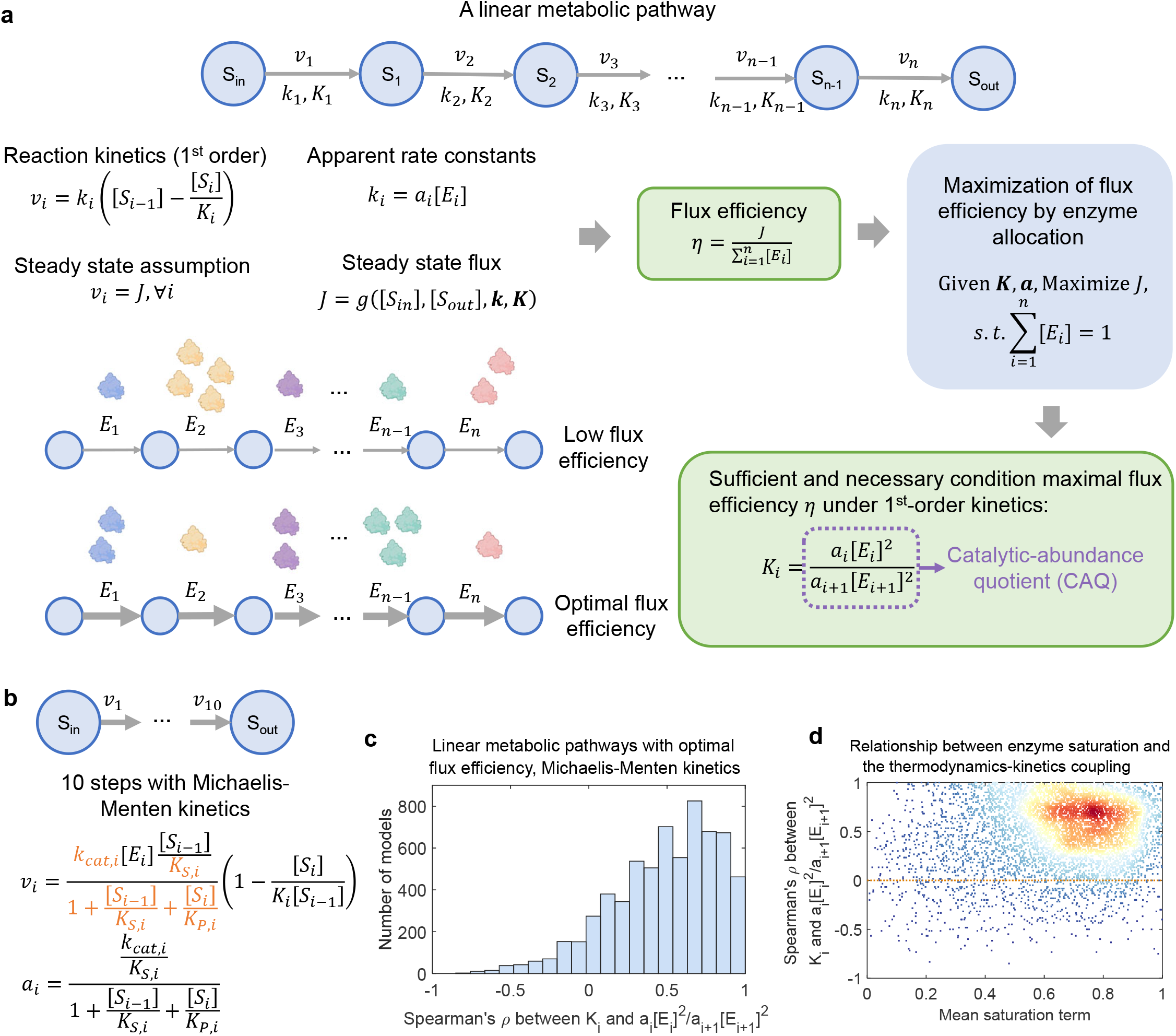
Law of kinetics-thermodynamics coupling underlying optimal efficiency of metabolic pathways. (a) The mathematical model for maximization of flux efficiency in a linear metabolic pathway with first-order kinetics. (b) Scheme of a linear metabolic pathway model with reversible Michaelis-Menten kinetics. (c) Bar plot showing the distribution of Spearman’s rank correlation coefficient between K and CAQ at optimal efficiency for the linear metabolic pathway with Michaelis-Menten kinetics. (d) Scatter plot comparing mean enzyme saturation term and Spearman’s rank correlation coefficient between K and CAQ at optimal efficiency for the linear metabolic pathway with Michaelis-Menten kinetics.

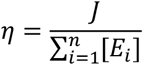

We then derived the conditions for the pathway to maximize the flux efficiency *η*. As the rate of each reaction is linear to its apparent rate constant *k*, which is proportional to the enzyme abundance [*E*], multiplying each enzyme abundance [*E*_*i*_] by a constant *α* while keeping [*S*_*in*_], [*S*_*out*_] and ***K*** unchanged does not affect the value of *η*:

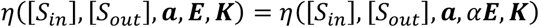

Therefore, without the loss of generality, the problem of maximizing the flux efficiency *η* is equivalent to the constrained optimization problem below:

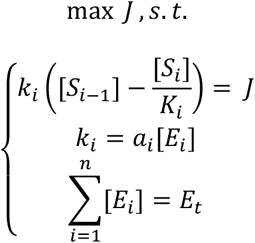

In which *E*_*t*_ is the total abundance of enzymes available to this pathway. Using Lagrange’s multiplier method (Supplementary Information), we can prove that the relationship below holds for every tandem reaction pair at maximal efficiency:

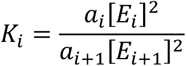

This law of thermodynamics-kinetics coupling states that the equilibrium constant of the upstream reaction, *K*_*i*_, must equal the ratio of the catalytic efficiencies multiplied by the squared enzyme abundance ratio, a term we define as the catalytic-abundance quotient (CAQ).

Next, we extended our theoretical analysis to zero-order kinetics and reversible Michaelis-Menten kinetics, deriving conditions for optimal pathway efficiency for both. For each of these reaction kinetics, the optimal condition has a generalized form (Fig S2a, Supplementary Information):

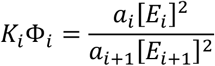

In which 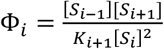 for zero-order kinetics and Φ_*i*_ = 1 for first-order kinetics. For Michaelis-Menten kinetics, the form of Φ _*i*_ depends on the concentrations of metabolites, equilibrium constants, and Michaelis constants related to the two reactions *r*_*i*_ and *r*_*i*+1_. Nevertheless, despite the complexity in the form of Φ _*i*_, the coupling between *K*_*i*_ and CAQ persists under Michaelis-Menten kinetics, as numerical simulation of a 10-reaction pathway (Fig. 2b, Methods) showed strong positive correlation between *K*_*i*_ and CAQ across almost all parameter configurations (Fig. 2c) independent of enzyme saturation levels (Fig. 2d, Fig. S2).

Finally, we extended the analysis to branching pathways (Fig. S3). Optimization of flux efficiency in these pathways can be solved by minimizing the total enzyme abundance under given steady state flux configuration, since it is impossible to define a single flux efficiency metric due to the coexistence of multiple branches. We considered both diverging and converging pathways (Fig. S3b). Although strict equality between K and CAQ no longer holds, the optimality solution still obeys inequalities with similar form: for the diverging pathway, the relationship 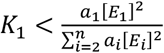 holds, while for the converging pathway, the relationship becomes 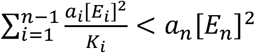 (Fig. S3b,c, Supplementary Information). Therefore, optimal enzyme allocation profiles in these pathways are still constrained by the relationship between K and CAQ.

To summarize, our theoretical analysis reveals the law of thermodynamics-kinetics coupling that links enzyme abundance, efficiency, and reaction thermodynamics at optimal efficiency of metabolic pathways. The mathematical simplicity of this law allows direct validation of it with omics data across diverse biological contexts.

### Validation of the law of thermodynamics-kinetics coupling

Validation of the law requires the equilibrium constant *K* of metabolic reactions and catalytic efficiency and abundance of metabolic enzymes. We estimated these parameters using a multi-step workflow (Fig. 3a, Methods). We computed the equilibrium constants from their standard Gibbs free energy changes, estimated catalytic efficiency of the enzymes from their *k*_*cat*_ and *K*_*m*_ values and substrate concentrations, and obtained enzyme abundances from proteomics datasets for 378 CCLE human cell lines^19^, *S. cerevisiae*^20^, and *E. coli*^21^. Moreover, acknowledging that the protein abundances and transcript levels are substantially correlated at cross-gene level^22, 23^ (Fig. S1), we also used transcriptomics datasets for human tumor samples and normal tissue samples in the TCGA project to validate our theory. These datasets were combined to compute the CAQ for each tandem reaction pair, which was finally correlated with the equilibrium constants of upstream reactions.

**Figure 3.**
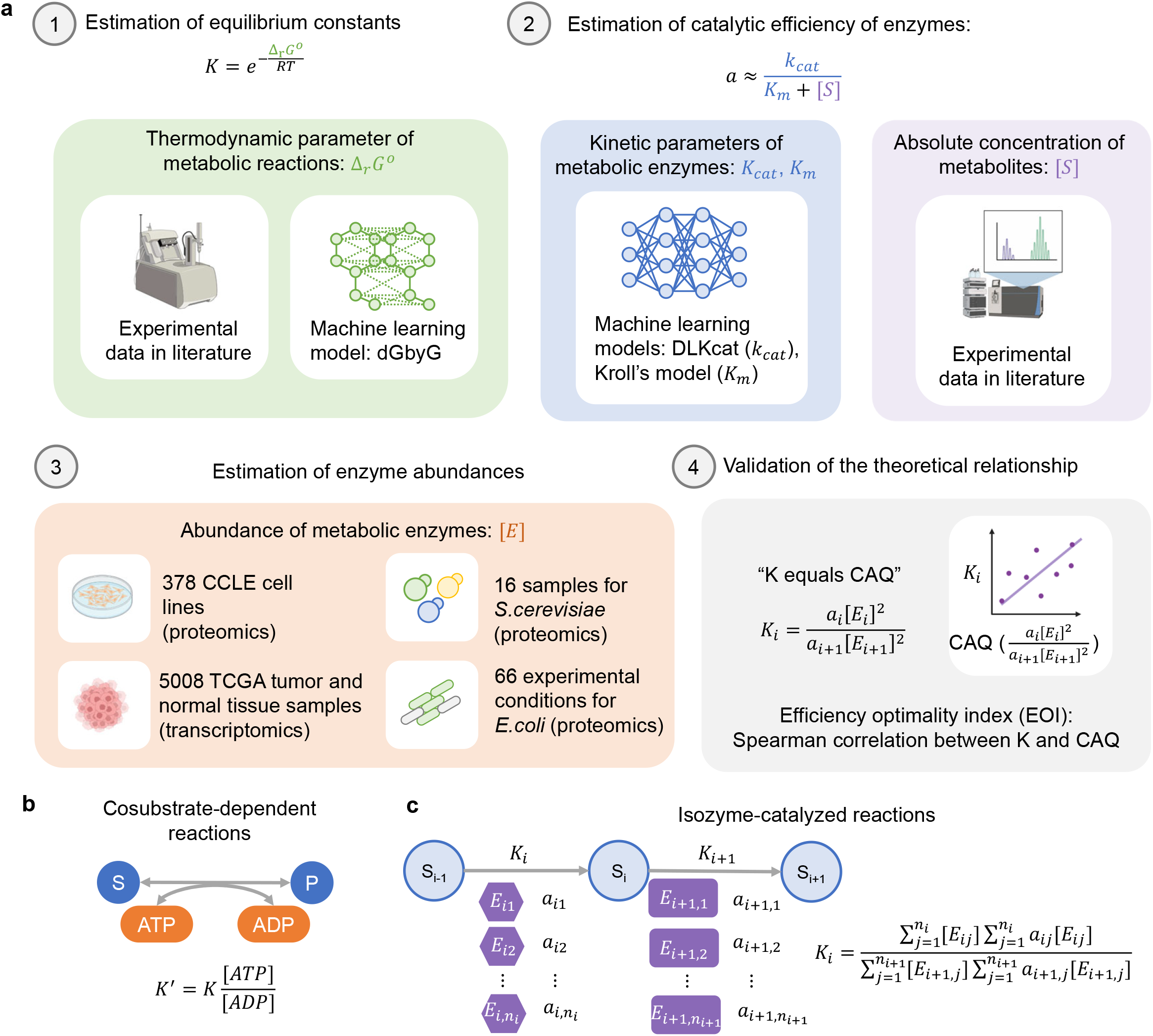
Validation of the law of thermodynamics-kinetics coupling. (a) Workflow for validation of the law with thermodynamics, enzyme kinetics, and omics datasets. (b) Adjustment of equilibrium constants for reactions depending on cofactors such as ATP/ADP. (c) Law of thermodynamics-kinetics coupling for reactions catalyzed by isozymes.

We next address two complexities of real-world metabolism that are not present in our initial theoretical analysis: the involvement of cofactors and isozymes. For reactions depending on cofactors (ATP/ADP, NADP^+^/NADPH, NAD^+^/NADH, and so on), we combined experimentally measured intracellular concentrations of these cofactors in literature to compute adjusted equilibrium constants *K*′ which were used for validation (Methods). For reactions catalyzed by isozymes, whose relative abundances are largely conserved across tissue types (Fig. S4), we treated their relative abundances as constant. Under this assumption, we derived a generalized form of the law, in which the CAQ term involves the catalytic efficiencies and abundances of all isozymes (Supplementary Information):

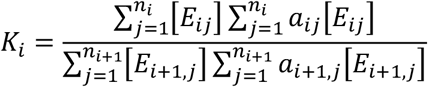

In which *a*_*i j*_ and [*E*_*i j*_] are catalytic efficiency and concentration of the *j*-th isozyme catalyzing the *i*-th reaction. This formulation preserves the core principle that K equals the CAQ, which is a quotient linking catalytic efficiencies and abundances of enzymes participating in two connected reactions.

Finally, we quantified the agreement between theory and data by calculating Spearman’s rank correlation coefficient between K and CAQ across all tandem reaction pairs in a pathway. We term this correlation the Efficiency Optimality Index (EOI). The theory predicts that at optimal efficiency, K equals CAQ for every pair, resulting in a perfect correlation (EOI = 1).

### Glycolysis obeys the law of thermodynamics-kinetics coupling across species

Glycolysis is an ancient pathway conserved across the kingdoms of life producing energy from carbohydrates. It is of particular interest to systems biologists because it is conserved across species, plays central role in carbon metabolism, and is subject to complex regulations^24^ that give rise to rich dynamic patterns such as oscillation^25^ and multistability^26^. Additionally, the well-characterized steps and enzymes in glycolysis (Fig. 4a), along with its near-linear structure and the availability of quantitative experimental data across biological contexts, makes it ideal for mathematical modeling. Here, we use omics and enzyme kinetics data from three phylogenetically distant species, *H. sapiens, S. cerevisiae*, and *E. coli* to validate the law of thermodynamics-kinetics coupling.

**Figure 4.**
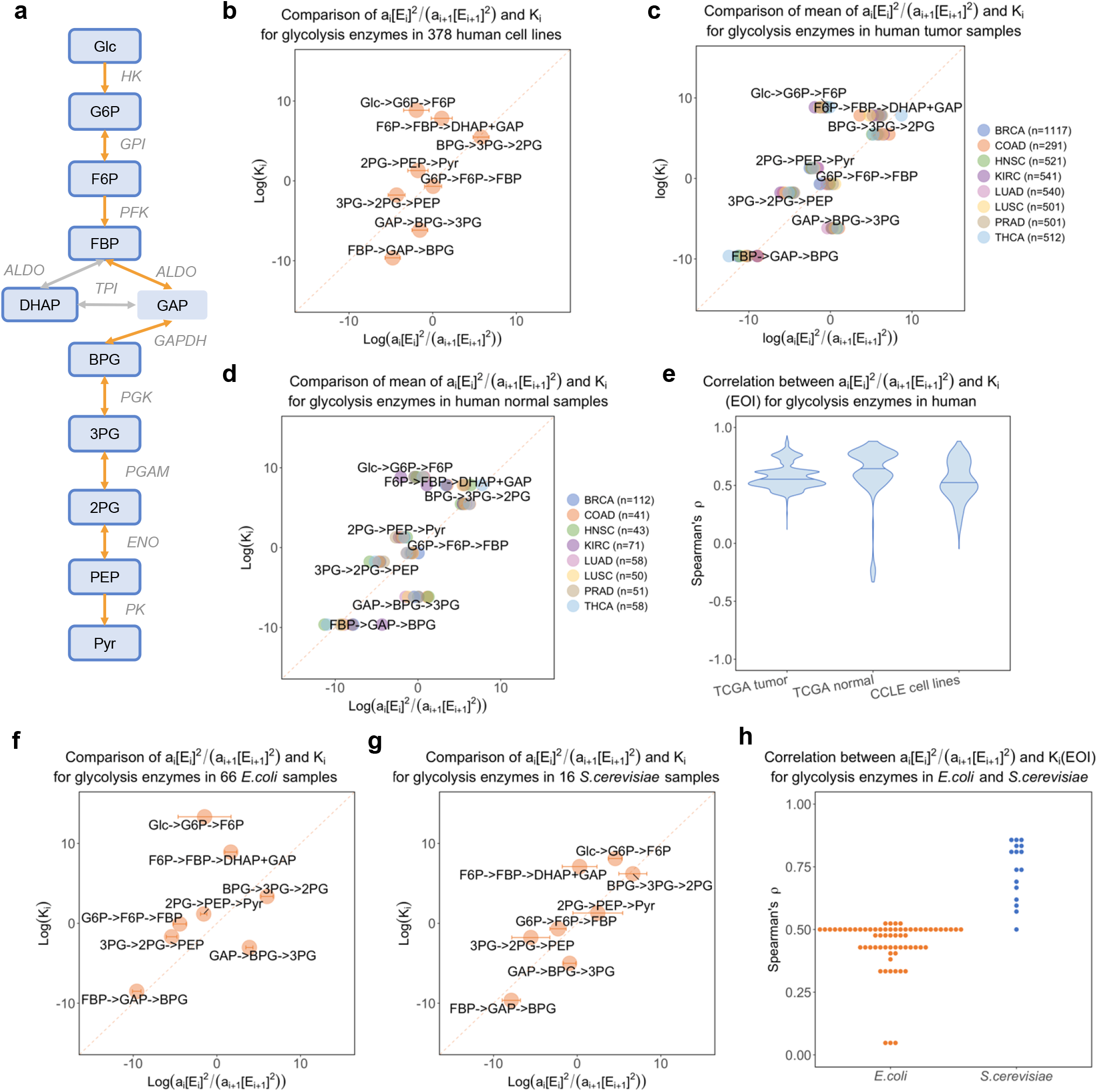
Glycolysis obeys the law of thermodynamics-kinetics coupling across species. (a) Diagram of the glycolysis pathway. (b) Scatter plot comparing K and CAQ (*a*_*i*_[*E*_*i*_]^2^/(*a*_*i*+1_[*E*_*i*+1_]^2^)) of tandem reaction pairs in glycolysis for 378 CCLE human cell lines. (c) Same as in (b) but for TCGA human tumor samples. (d) Same as in (b) but for human normal tissue samples in TCGA. (e) Distribution of Spearman’s rank correlation coefficient between K and CAQ, i.e. EOI of glycolysis, for the human cell lines, tumors, and normal tissue samples. (f) Same as in (b) but for the bacteria *Escherichia coli*. (g) Same as in (b) but for the budding yeast *Saccharomyces cerevisiae*. (h) Distribution of the EOI of glycolysis in the *E*.*coli* and *S*.*cerevisiae* samples.

We first retrieved proteomics data for 378 human cell lines from the CCLE database and transcriptomics data for 5008 human tumor and normal tissue samples from the TCGA project covering eight different tissues-of-origin, including breast invasive carcinoma (BRCA), colon adenocarcinoma (COAD), head and neck squamous cell carcinoma (HNSC), kidney renal clear cell carcinoma (KIRC), lung adenocarcinoma (LUAD), lung squamous cell carcinoma (LUSC), prostate adenocarcinoma (PRAD), and thyroid carcinoma (THCA). We computed the CAQ of all tandem reaction pairs in glycolysis, compared them to the upstream equilibrium constants (*K*_*i*_) (Fig. 4b-d), and computed the EOI of glycolysis by correlating these two for each sample. The results reveal a robust coupling between K and CAQ across human cell and tissue types (median EOI > 0.5 for both TCGA samples and CCLE cell lines, Fig. 4e), consistent with the theory. This coupling relationship was conserved in the unicellular organisms *S. cerevisiae* (Fig. 4f) and *E. coli* (Fig. 4g) across diverse experimental conditions (Fig. 4h) and remained unchanged when Pearson’s correlation coefficients were used to assess their correlation (Fig. S5). Lower EOI values were observed if substrate concentrations were not used in estimating the catalytic efficiencies (Methods), highlighting the importance of quantitative metabolite concentrations in validation of the theory (Fig. S6). These results demonstrate that glycolysis follows the law of thermodynamics-kinetics coupling in a variety of cell types covering both human and unicellular organisms.

Next, we investigated potential mechanisms enabling the optimal efficiency state. There are two strategies to establish the coupling between K and CAQ: (1) by tuning enzyme abundance [*E*] under a given set of catalytic efficiencies *a*, or (2) by evolving enzyme kinetic parameters such as *k*_*cat*_ and *K*_*m*_ (thus the corresponding *a*) so that the thermodynamics-kinetics coupling is robust to variations in [*E*]. Since evolution of DNA and protein sequences can change both expression level (strategy 1) and kinetic parameters (strategy 2) of an enzyme, optimization of flux efficiency can involve both strategies. To characterize their contributions, we performed permutation tests in which the catalytic efficiencies and abundances of glycolytic enzymes were individually or jointly permuted. If the cell relies on strategy 1 only, random permutation of enzyme abundance will completely impair the correlation between *K* and CAQ. Otherwise, the correlation between K and CAQ will persist after permutation of the enzyme abundances. Disrupting enzyme abundances alone preserved a significantly positive EOI in all human (Fig. S7a-c), bacteria (Fig. S7d) and yeast (Fig. S7e) datasets, while the coupling diminished when both enzyme abundances and catalytic efficiencies were permuted. Therefore, evolution of metabolic pathways likely utilizes both strategies: enzyme kinetic parameters are pre-optimized to provide robustness (strategy 1), and cells further fine-tune enzyme expression (strategy 2) to achieve optimal efficiency.

Taking together, these results suggest that the enzyme allocation of glycolysis robustly obeys the law of thermodynamics-kinetics coupling across species, although evolutionary separation of the three species from their common ancestors occurred billions of years ago. This evolutionary convergence highlights the fundamental role of efficient resource utilization in shaping metabolism.

### Determinants of glycolytic efficiency optimization

While the thermodynamics-kinetics coupling was robustly observed across species, the resulting EOI still varied across cell types (Fig. 4e) and experimental conditions (Fig. 4h). Therefore, we sought to identify potential determinants of the EOI of glycolysis. We correlated gene expression levels to EOI of glycolysis in each type of TCGA tumors, and performed gene set enrichment analysis (GSEA) to identify KEGG pathways associated with higher EOI in each cancer type (Fig. 5a). We found that higher glycolytic EOI was consistently associated with the activation of multiple oncogenic signaling pathways across most cancer types (Fig. 5b,c, Fig S8), suggesting that aggressive tumors have stronger drive to optimize glycolytic efficiency. Another universal trend observed in the TCGA tumors is negative correlation between the EOI of glycolysis and the expression of ribosome (Fig. S9a,b) and oxidative phosphorylation (OXPHOS) genes (Fig. S9c,d). This trend indicates a potential trade-off between glycolytic efficiency and other energy-consuming processes and alternative energy-producing routes.

**Figure 5.**
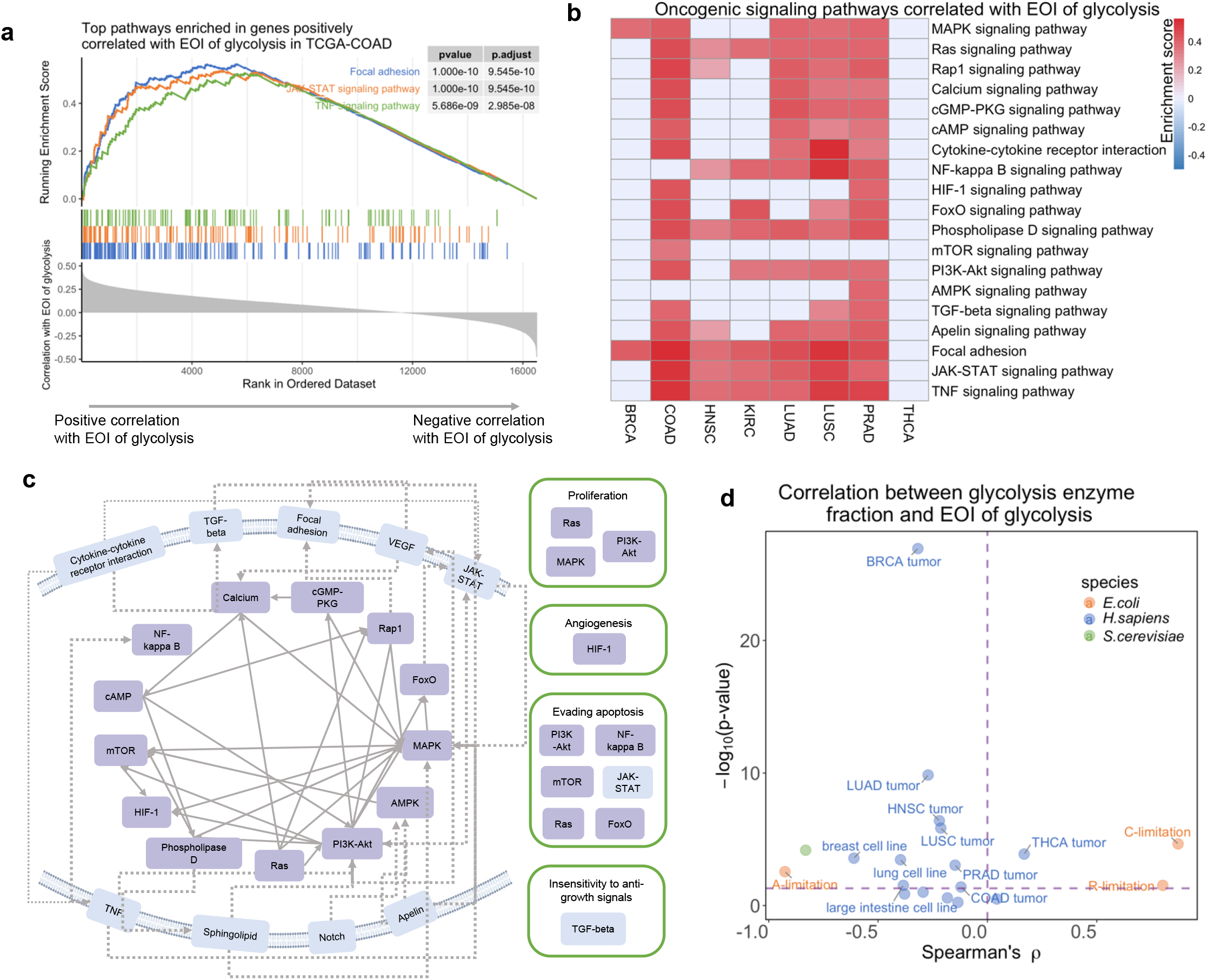
Determinants of glycolytic efficiency optimization. (a) Top pathways enriched in genes positively correlated with EOI of glycolysis in TCGA-COAD. (b) Enrichment scores for oncogenic signaling pathways positively associated with EOI of glycolysis. (c) Regulatory links between oncogenic signaling pathways associated with EOI of glycolysis. (d) Spearman’s rank correlation between fraction of glycolytic enzyme in total cellular proteome and EOI of glycolysis.

Another factor that may affect the optimization of glycolytic efficiency is the total cellular investment in glycolytic enzymes. We computed the fraction of glycolytic enzymes in the total cellular proteome and correlated it to the EOI of glycolysis. A negative correlation between these two was observed in most of the samples (15 negative correlations out of 19 groups of samples, Fig. 5d, Fig. S10), suggesting that a tighter constraint on total available glycolytic enzyme abundance is a key driver for maximization of the pathway efficiency. This mirrors a principle of rational economics in which optimal efficiency was necessitated under scarcity of resources.

Taken together, these results highlight the heterogeneous nature of glycolytic efficiency optimization and reveal potential determinants of it. The link to oncogenic signaling suggests that pathological processes could change the economic decision of cells, while the inverse relationship with proteomic investment provides clue that resource constraints may shape metabolic efficiency.

## Discussion

In this study, we derived the law of thermodynamics-kinetics coupling underlying optimal proteome efficiency of metabolic pathways, and validated it with thermodynamics, kinetics, and omics datasets across species and cell types, demonstrating that efficiency optimization is a fundamental principle operating from microbes to human cells. The simplicity of this law opens a route for rationally engineering synthetic pathways towards maximal efficiency, addressing a key challenge where engineered metabolic pathways often suffer from extremely low efficiency due to the lack of a predictive theoretical framework.

Our work advances the field of metabolic optimality in several key aspects. First, unlike previous theoretical studies that rely on complex numerical solutions (e.g. the ECM method^17^), our work establishes a simple, general law readily applicable to realistic metabolic pathways with biological complexities such as cofactors and isozymes. Numerical simulations confirm its robustness across different rate laws, in which the core coupling between K and CAQ holds under the most complex reversible Michaelis-Menten kinetics. Although realistic metabolic enzymes are usually substrate-saturated^27^, the successful validation of the law with glycolytic enzymes across species supports its generalization to various kinetic regimes.

Second, our theory addresses a distinct level of proteome optimization. While previous coarse-grained models focus on global allocation among major functional sectors such as ribosome and metabolism^9-11^, our theory operates at local pathway level, determining how enzymes are optimally distributed among reaction steps to maximize the efficiency of each pathway. Hence the distinction parallels that between macro-and micro-economics, together illustrating how economic principles operate at multiple scales to achieve optimal cellular resource allocation.

Another important output of our study is the EOI metric that assesses how closely a pathway adheres to the theoretical optimum of efficiency. This enables systematic investigation of efficiency optimality across biological conditions. Applying this metric, we found that two key patterns emerge: first, higher EOI was associated with oncogenic signaling in human tumors, suggesting that malignant cells not only upregulate but also optimize glycolysis; second, EOI was negatively correlated with total proteomic investment in glycolysis, revealing a “scarcity-driven efficiency” where tighter resource constraints necessitate stronger efficiency optimization. This pattern is reminiscent of human rational decision-making under budget limitations, indicating an intrinsic similarity between fundamental principles of natural and societal processes.

Looking forward, extending the theory and validation to genome-scale metabolic networks (GEMs) is promising yet challenging due to the highly nonlinear topology of metabolic networks and the current scarcity of absolute metabolite concentration data required for accurate estimation of catalytic efficiencies. It is also important to determine how regulatory links within metabolic networks affect the optimal allocation of enzymes. However, with our theoretical work on efficiency optimality of branching pathways and as metabolomics technologies advance, we anticipate that large-scale testing of the thermodynamics-kinetics law will become feasible, further illuminating the economic principles that govern cellular metabolism at a systems level.

Furthermore, the variation in EOI across conditions highlights that proteome efficiency is one of several competing metabolic objectives. Other critical goals such as robustness to environmental fluctuations^28, 29^ and minimization of metabolite load also have profound effects, resulting in pre-optimized thermodynamic pattern of pathways^30^, precise tuning of metabolite concentrations^31^ and scaling between metabolites and enzymes^32^. A complete understanding of how cells balance these objectives requires both theoretical derivation of the Pareto optimal solutions and integration of multi-omics data to map how different cellular states navigate this Pareto landscape. Future work precisely defining this multi-objective Pareto front will be essential for a systematic understanding of metabolic design principles.

## Methods

### Linear pathway model with Michaelis-Menten kinetics

The mathematical model of a linear metabolic pathway consisting of ten reactions with reversible Michaelis-Menten kinetics was implemented with MATLAB. Reaction rate of the i-th reaction in the pathway is calculated as follows:

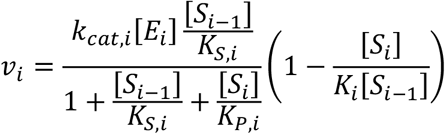

Therefore, the dynamics of the metabolite concentrations in the pathway can be modeled using the ordinary differential equations (ODEs) below:

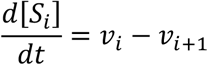

The reaction thermodynamic and kinetic parameters and boundary metabolite concentrations were randomly sampled from log-normal distributions:

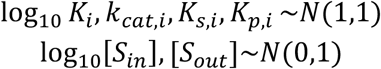

We randomly generated 10,000 parameter sets from the distributions. Only parameter sets producing positive pathway flux were kept for the following analyses. For each parameter set and a given set of enzyme concentrations, the steady state concentrations of intermediate metabolites [*S*_*i*_] was computed by solving the steady state equations:

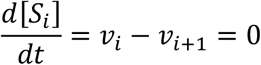

This problem was solved by minimizing the squared sum of all *v*_*i*_ − *v*_*i*+1_ terms using the MATLAB function lsqnonlin[] to address numerical difficulties of directly solving the nonlinear equations. Steady state pathway flux was then computed from metabolite concentrations. Optimal enzyme allocation profile maximizing flux efficiency under a given parameter set was computed using the MATLAB constrained optimization solver fmincon[] to maximize the pathway flux with the upper bound of total enzyme abundance set to 1. Saturation term of the i-th step was computed as follows:

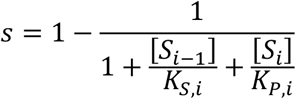

### Estimation of equilibrium constant

Equilibrium constants (*K*) of each reaction were calculated from standard Gibbs energy changes (Δ_*r*_*G*^*o*^) based on the equation Δ_*r*_*G*^*o*^ = −*RT* ln *K*, with R=8.314472 J·mol^− 1^·K^− 1^, T=298.15K (*H*.*sapiens* and *S*.*cerevisiae*), or T=310.15K (*E*.*coli*). Standard Gibbs free energy changes(ΔG°’) of glycolysis reactions for eukaryotes were obtained from experimental data in literature^33^. For *E*.*coli*, ΔG°’ values were predicted using the machine learning model dGbyG^30^, under physiological conditions compiled from literature^34, 35^ (Dataset S1). Subsequently, the adjusted equilibrium constant (*K*′) were calculated from the original equilibrium constant K and intracellular concentrations of cofactors by multiplying the ratio of the product of concentrations of cofactors serving as products to the product of concentrations of cofactors serving as substrates to the original equilibrium constant K. For example, the phosphorylation of glucose catalyzed by hexokinase has the chemical equation below:

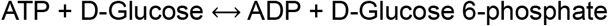

Here we treated ATP and ADP as cofactors and used their intracellular concentrations to compute the adjusted equilibrium constant:

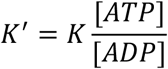

The adjusted equilibrium constants in the three organisms are available in Dataset S2. Substrate concentrations were obtained from published datasets^36-39^ (Dataset S3).

### Estimation of catalytic efficiency

We estimated the catalytic efficiency of metabolic enzymes using reversible Michaelis-Menten kinetics considering substrate saturation. In this case, the catalytic efficiency of an enzyme can be computed from its kinetic parameters *k*_*cat*_ and *K*_*m*_, together with the concentration of substrate:

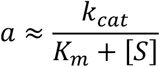

For the enzyme kinetic parameters *k*_*cat*_ and *K*_*m*_, since experimental values from the BRENDA enzyme database^40^ are missing for the wildtype form of several enzymes in glycolysis and individual measurements from different sources vary a lot, we used values predicted based on amino acid sequences of the catalytic subunit of each enzyme and molecular structures of its substrates by two deep learning algorithms, including *k*_*cat*_ values predicted by DLKcat^41, 42^ and *K*_*m*_ values predicted by the machine learning model developed by Kroll *et al*^43^. Amino acid sequences of the enzymes obtained from UniProt and SMILES strings encoding structures of the metabolites retrieved from MetaNetX^44^ were used in the prediction^45^. The predicted values of enzyme kinetic parameters were available in Dataset S4. Experimentally measured absolute metabolite concentrations^27, 38, 46, 47^ in human cells, the bacteria *Escherichia coli*, and the unicellular eukaryote *Saccharomyces cerevisiae* were used together with the enzymatic parameters to estimate the catalytic efficiency *a*.

A specific case that needs to be addressed is the GLCptspp reaction catalyzed by the phosphotransferase system (PTS) in *E. coli*. The PTS consists of three components, enzyme I (EI), histidine phosphocarrier protein (HPr), and membrane bounded sugar specific permeases (enzymes II, EII)^48^. In the iML1515 model, the gene-reaction rules for GLCptspp include the following genes: *ptsH* (EI), *ptsI* (HPr), *crr* (EII), *ptsG* (EII), *malX* (EII), *manX* (EII), *manY* (EII), and *manZ* (EII). We used amino acid sequence of the subunit ptsG responsible for specific glucose binding and catalysis (EC:2.7.1.199) in prediction of the kinetic parameters.

### Omics data processing

Information about reactions in glycolysis, including their substrates, products and gene-protein-reaction relationships, were extracted from the following genome-scale metabolic models (GEMs): Recon3D (*H*.*sapiens*)^49^, iML1515 (*E*.*coli*)^50^, and iMM904 (*S*.*cerevisiae*)^51^. This study utilized three published proteome datasets. The proteomics data for 378 human cancer cell lines was obtained from the Cancer Cell Line Encyclopedia (https://sites.broadinstitute.org/ccle/datasets)^52^. The mass spectrometry intensities of the proteins were normalized into molar fractions using molecular mass of the enzymes. The proteomics datasets for *E*.*coli* and *S*.*cerevisiae* were retrieved from two published datasets summarizing absolute protein copy numbers in the two species across independent proteomics studies^53, 54^.

Transcriptomics datasets in human tumors and normal tissue samples were obtained from The Cancer Genome Atlas (TCGA) database (https://www.cancer.gov/tcga). Transcripts Per Million (TPM) data for BRCA, HNSC, KIRC, LUAD, LUSC, PRAD, and THCA were downloaded directly. Original paired-end FASTQ files for COAD were downloaded and mapped to the human genome (GRCh38) using STAR (version 2.7.9a)^55^. Read counts were then calculated using featureCounts (version 1.5.3)^56^, and subsequently normalized into TPM using R (version 4.3.2). Zero expression levels were replaced with a small positive number of 10^−6^ to avoid zero or infinity values of the resulting CAQ. Fraction of glycolytic enzymes in the entire cellular proteome was computed by dividing total protein copy number of glycolytic enzymes by the total copy number of all proteins (proteomics datasets) or approximated by computing total TPM of all glycolytic genes (transcriptomics data).

### Gene set enrichment analysis (GSEA)

All transcriptome datasets were divided into tumor and normal groups based on sample information. Within each group, the mean TPM value for each gene was calculated for samples sharing the same EOI value. Spearman’s correlation coefficient between each gene’s expression values and EOI was then computed, representing the degree to which the gene’s expression is coupled with glycolytic efficiency. Genes were ranked based on this coefficient. KEGG pathway enrichment analysis was performed in R (version 4.3.2) using the clusterProfiler package (version 4.10.1)^57^ with an adjusted p-value (*P*_*adj*_) cutoff of <0.05.

## Data availability

Thermodynamics, kinetics, and omics data used in this study are available at the Github repository: https://github.com/bioinf-kud/Optimality-in-glycolysis-enzyme-allocation.

## Code availability

Scripts for validation of the theory are available at the Github repository: https://github.com/bioinf-kud/Optimality-in-glycolysis-enzyme-allocation. Scripts for numerical simulation of the linear metabolic pathway model with Michaelis-Menten kinetics are available at the Github repository: https://github.com/ziweidai/Linear_metabolic_pathway_MM_kinetics.

## Acknowledgements

The authors thank all members of the Dai lab for helpful discussions. We thank the National Natural Science Foundation of China (12371489 to ZD), National Key Research and Development Program of China (2021YFA0911300 to ZD), Guangdong Program (2021QN02Y856 to ZD), and the Shenzhen Science and Technology Program (KQTD20180411143432337 to ZD) for their generous support.

## Author contributions

K.S., W.Z. and Z.D. designed the study and wrote the manuscript. Z.D. performed all mathematical derivations and numerical simulation of the linear pathway model. K.S. and W.Z. analyzed the omics datasets with inputs from D.H. and conducted all validation of the theoretical results. W.F. predicted the standard Gibbs free energy of *E*.*coli* reactions. K.S. and C.D. performed the GSEA analyses. All authors have read and approved the final manuscript.

## References

1. Islam, M.M. et al. Reframing the role of the objective function in its proper context for metabolic network modeling. Cell Syst 16, 101298 (2025).

2. Welle, S. & Nair, K.S. Relationship of resting metabolic rate to body composition and protein turnover. Am J Physiol 258, E990–998 (1990).

3. Akashi, H. & Gojobori, T. Metabolic efficiency and amino acid composition in the proteomes of Escherichia coli and Bacillus subtilis. Proc Natl Acad Sci U S A 99, 3695–3700 (2002).

4. Heizer, E.M., Jr. et al. Amino acid cost and codon-usage biases in 6 prokaryotic genomes: a whole-genome analysis. Mol Biol Evol 23, 1670–1680 (2006).

5. Chen, Y. & Nielsen, J. Yeast has evolved to minimize protein resource cost for synthesizing amino acids. Proc Natl Acad Sci U S A 119 (2022).

6. Zhang, H. et al. Biosynthetic energy cost for amino acids decreases in cancer evolution. Nat Commun 9, 4124 (2018).

7. Piper, M.D.W. et al. Matching Dietary Amino Acid Balance to the In Silico-Translated Exome Optimizes Growth and Reproduction without Cost to Lifespan. Cell Metab 25, 1206 (2017).

8. Shenhav, L. & Zeevi, D. Resource conservation manifests in the genetic code. Science 370, 683–687 (2020).

9. Scott, M., Gunderson, C.W., Mateescu, E.M., Zhang, Z. & Hwa, T. Interdependence of cell growth and gene expression: origins and consequences. Science 330, 1099–1102 (2010).

10. Basan, M. et al. Overflow metabolism in Escherichia coli results from efficient proteome allocation. Nature 528, 99–104 (2015).

11. Zhu, M., Mori, M., Hwa, T. & Dai, X. Distantly related bacteria share a rigid proteome allocation strategy with flexible enzyme kinetics. Proc Natl Acad Sci U S A 122, e2427091122 (2025).

12. Flamholz, A.I., Goyal, A., Fischer, W.W., Newman, D.K. & Phillips, R. The proteome is a terminal electron acceptor. Proc Natl Acad Sci U S A 122, e2404048121 (2025).

13. Khana, D.B. et al. Thermodynamics shapes the in vivo enzyme burden of glycolytic pathways. mBio, e0183725 (2025).

14. Heinrich, R. & Klipp, E. Control analysis of unbranched enzymatic chains in states of maximal activity. J Theor Biol 182, 243–252 (1996).

15. Brown, G.C. Total cell protein concentration as an evolutionary constraint on the metabolic control distribution in cells. J Theor Biol 153, 195–203 (1991).

16. Klipp, E. & Heinrich, R. Competition for enzymes in metabolic pathways: implications for optimal distributions of enzyme concentrations and for the distribution of flux control. Biosystems 54, 1–14 (1999).

17. Noor, E. et al. The Protein Cost of Metabolic Fluxes: Prediction from Enzymatic Rate Laws and Cost Minimization. PLoS Comput Biol 12, e1005167 (2016).

18. Noor, E. & Liebermeister, W. Optimal enzyme profiles in unbranched metabolic pathways. Interface Focus 14, 20230029 (2024).

19. Nusinow, D.P. et al. Quantitative Proteomics of the Cancer Cell Line Encyclopedia. Cell 180, 387–402 e316 (2020).

20. Ho, B., Baryshnikova, A. & Brown, G.W. Unification of Protein Abundance Datasets Yields a Quantitative Saccharomyces cerevisiae Proteome. Cell Syst 6, 192–205 e193 (2018).

21. Mori, M. et al. From coarse to fine: the absolute Escherichia coli proteome under diverse growth conditions. Mol Syst Biol 17, e9536 (2021).

22. Buccitelli, C. & Selbach, M. mRNAs, proteins and the emerging principles of gene expression control. Nat Rev Genet 21, 630–644 (2020).

23. Schwanhausser, B. et al. Global quantification of mammalian gene expression control. Nature 473, 337–342 (2011).

24. Nemkov, T. et al. Biological and genetic determinants of glycolysis: Phosphofructokinase isoforms boost energy status of stored red blood cells and transfusion outcomes. Cell Metab 36, 1979–1997 e1913 (2024).

25. Chandra, F.A., Buzi, G. & Doyle, J.C. Glycolytic oscillations and limits on robust efficiency. Science 333, 187–192 (2011).

26. Mulukutla, B.C., Yongky, A., Daoutidis, P. & Hu, W.-S. Bistability in Glycolysis Pathway as a Physiological Switch in Energy Metabolism. PLOS ONE 9, e98756 (2014).

27. Park, J.O. et al. Metabolite concentrations, fluxes and free energies imply efficient enzyme usage. Nat Chem Biol 12, 482–489 (2016).

28. Shen, Y. et al. Mitochondrial ATP generation is more proteome efficient than glycolysis. Nat Chem Biol 20, 1123–1132 (2024).

29. Basan, M. et al. A universal trade-off between growth and lag in fluctuating environments. Nature 584, 470–474 (2020).

30. Fan, W. et al. Unraveling principles of thermodynamics for genome-scale metabolic networks using graph neural networks. Cell Syst, 101393 (2025).

31. Tepper, N. et al. Steady-state metabolite concentrations reflect a balance between maximizing enzyme efficiency and minimizing total metabolite load. PLoS One 8, e75370 (2013).

32. Dourado, H., Mori, M., Hwa, T. & Lercher, M.J. On the optimality of the enzyme-substrate relationship in bacteria. PLoS Biol 19, e3001416 (2021).

33. Mathews, C.K. Biochemistry, Edn. 4th ed. (Pearson, Toronto; 2013).

34. Hurwitz, C. & Rosano, C.L. The Intracellular Concentration of Bound and Unbound Magnesium Ions in Escherichia coli. Journal of Biological Chemistry 242, 3719–3722 (1967).

35. Noor, E., Haraldsdóttir, H.S., Milo, R. & Fleming, R.M.T. Consistent Estimation of Gibbs Energy Using Component Contributions. PLOS Computational Biology 9, e1003098 (2013).

36. Park, J.O. et al. Metabolite concentrations, fluxes and free energies imply efficient enzyme usage. Nature Chemical Biology 12, 482–489 (2016).

37. Thorfinnsdottir, L.B., García-Calvo, L., Bø, G.H., Bruheim, P. & Røst, L.M. Optimized Fast Filtration-Based Sampling and Extraction Enables Precise and Absolute Quantification of the Escherichia coli Central Carbon Metabolome. Metabolites 13, 150 (2023).

38. Bashir, A., Gropler, R. & Ackerman, J. Absolute Quantification of Human Liver Phosphorus-Containing Metabolites In Vivo Using an Inhomogeneous Spoiling Magnetic Field Gradient. PLoS One 10, e0143239 (2015).

39. Chowdhury, H.H. Differences in cytosolic glucose dynamics in astrocytes and adipocytes measured by FRET-based nanosensors. Biophysical Chemistry 261, 106377 (2020).

40. Chang, A. et al. BRENDA, the ELIXIR core data resource in 2021: new developments and updates. Nucleic Acids Res 49, D498–D508 (2021).

41. Li, F. et al. Deep learning-based kcat prediction enables improved enzyme-constrained model reconstruction. Nature Catalysis 5, 662–672 (2022).

42. Li, F., Chen, Y., Anton, M. & Nielsen, J. GotEnzymes: an extensive database of enzyme parameter predictions. Nucleic Acids Res 51, D583–D586 (2023).

43. Kroll, A., Engqvist, M.K.M., Heckmann, D. & Lercher, M.J. Deep learning allows genome-scale prediction of Michaelis constants from structural features. PLoS Biol 19, e3001402 (2021).

44. Moretti, S., Tran, V.D.T., Mehl, F., Ibberson, M. & Pagni, M. MetaNetX/MNXref: unified namespace for metabolites and biochemical reactions in the context of metabolic models. Nucleic Acids Research 49, D570–D574 (2021).

45. Consortium, T.U. UniProt: the Universal Protein Knowledgebase in 2025. Nucleic Acids Research 53, D609–D617 (2025).

46. Chowdhury, H.H. Differences in cytosolic glucose dynamics in astrocytes and adipocytes measured by FRET-based nanosensors. Biophys Chem 261, 106377 (2020).

47. Thorfinnsdottir, L.B., Garcia-Calvo, L., Bo, G.H., Bruheim, P. & Rost, L.M. Optimized Fast Filtration-Based Sampling and Extraction Enables Precise and Absolute Quantification of the Escherichia coli Central Carbon Metabolome. Metabolites 13 (2023).

48. Kotrba, P., Inui, M. & Yukawa, H. Bacterial phosphotransferase system (PTS) in carbohydrate uptake and control of carbon metabolism. Journal of Bioscience and Bioengineering 92, 502–517 (2001).

49. Brunk, E. et al. Recon3D enables a three-dimensional view of gene variation in human metabolism. Nature Biotechnology 36, 272–281 (2018).

50. Monk, J.M. et al. iML1515, a knowledgebase that computes Escherichia coli traits. Nature Biotechnology 35, 904–908 (2017).

51. Mo, M.L., Palsson, B.Ø. & Herrgård, M.J. Connecting extracellular metabolomic measurements to intracellular flux states in yeast. BMC Systems Biology 3, 37 (2009).

52. Nusinow, D.P. et al. Quantitative Proteomics of the Cancer Cell Line Encyclopedia. Cell 180, 387–402.e316 (2020).

53. Ho, B., Baryshnikova, A. & Brown, G.W. Unification of Protein Abundance Datasets Yields a Quantitative Saccharomyces cerevisiae Proteome. Cell Systems 6, 192–205.e193 (2018).

54. Mori, M. et al. From coarse to fine: the absolute Escherichia coli proteome under diverse growth conditions. Molecular Systems Biology 17, 1–23 (2021).

55. Dobin, A. et al. STAR: ultrafast universal RNA-seq aligner. Bioinformatics 29, 15–21 (2013).

56. Liao, Y., Smyth, G.K. & Shi, W. featureCounts: an efficient general purpose program for assigning sequence reads to genomic features. Bioinformatics 30, 923–930 (2014).

57. Wu, T. et al. clusterProfiler 4.0: A universal enrichment tool for interpreting omics data. The Innovation 2, 100141 (2021).

