## Supplementary Text for "A universal law of thermodynamics-kinetics coupling shapes enzyme allocation and glycolytic efficiency across species"

September 25, 2025

#### 1 Mathematical model of linear metabolic pathway

##### 1.1 Structure and thermodynamics of the linear pathway

We consider the simplest topology of a metabolic pathway, that is, a linear chain of reversible reactions. The structure of the pathway is shown below:

$$S_{in} \underset{}{\overset{k_1,K_1}{\rightleftarrows}} S_1 \underset{}{\overset{k_2,K_2}{\rightleftarrows}} S_2 \underset{}{\overset{k_3,K_3}{\rightleftarrows}} S_3 \cdots\cdots S_{n-1} \underset{}{\overset{k_n,K_n}{\rightleftarrows}} S_{out} \tag{1}$$

The rate of each reaction in the metabolic pathway is determined by first-order or zero-order kinetics, which we will define next. For both first-order and zero-order kinetics, parameters for one reaction in the pathway include an apparent rate constant $k$ and an equilibrium constant $K$. The equilibrium constant $K$ is linked to the standard Gibbs free energy change of this reaction, $\Delta_r G^o$:

$$\Delta_r G^o = -RT\log K \tag{2}$$

For a reaction with substrate $S$ and product $P$, its reaction Gibbs free energy $\Delta_r G$ is determined by $\Delta_r G^o$ and concentrations of the substrate and product:

$$\Delta_r G = \Delta_r G^o + RT\log\frac{[P]}{[S]} = RT\log\frac{[P]}{K[S]} \tag{3}$$

To simplfy the expression, let $g = \frac{\Delta_r G}{RT}$, thus we have:

$$g = \log\frac{[P]}{K[S]} \tag{4}$$

The reaction takes place in the forward direction when $g < 0$, backward direction when $g > 0$, and reaches thermodynamic equilibrium when $g = 0$. We refer to $g$ as the scaled Gibbs free energy change of the corresponding reaction as it differs from the reaction Gibbs free energy change $\Delta_r G$ by the scaling factor $\frac{1}{RT}$.

#### 1.2 First-order and zero-order kinetic laws

For the $i$-th reaction in this pathway, its reaction rate under first-order kinetics is:

$$v_i = k_i([S_i] - \frac{[S_{i+1}]}{K_i}) \tag{5}$$

Otherwise, the reaction rate under zero-order kinetics is:

$$v_i = k_i(1 - \frac{[S_{i+1}]}{K_i[S_i]}) \tag{6}$$

#### 1.3 Steady state solution of pathway flux

##### 1.3.1 First-order kinetics

We first sought to solve the steady state flux $J$ as a function of the input metabolite concentration $[S_{in}]$, the output metabolite concentration $[S_{out}]$, the vector of kinetic parameters $\mathbf{k}$, and the vector of equilibrium constants, $\mathbf{K}$. The pathway flux $J$ can be computed by solving the steady state equations:

$$\begin{cases} J = k_1([S_{in}] - \frac{[S_1]}{K_1}) \\ J = k_2([S_1] - \frac{[S_2]}{K_2}) \\ \quad\vdots \\ J = k_i([S_{i-1}] - \frac{[S_i]}{K_i}) \\ \quad\vdots \\ J = k_n([S_{n-1}] - \frac{[S_{out}]}{K_n}) \end{cases} \tag{7}$$

Multiplying a factor $\frac{1}{k_i \prod_{j=1}^{i-1} K_j}$ to the $i$-th equation, then we have:

$$\begin{cases} \frac{J}{k_1} = [S_{in}] - \frac{[S_1]}{K_1} \\ \frac{J}{k_2 K_1} = \frac{[S_1]}{K_1} - \frac{[S_2]}{K_1 K_2} \\ \quad\vdots \\ \frac{J}{k_i \prod_{j=1}^{i-1} K_j} = \frac{[S_{i-1}]}{\prod_{j=1}^{i-1} K_j} - \frac{[S_i]}{\prod_{j=1}^{i} K_j} \\ \quad\vdots \\ \frac{J}{k_n \prod_{j=1}^{n-1} K_j} = \frac{[S_{n-1}]}{\prod_{j=1}^{n-1} K_j} - \frac{[S_{out}]}{\prod_{j=1}^{n} K_j} \end{cases} \tag{8}$$

Summing up all equations, we have:

$$\left(\sum_{i=1}^{n} \frac{1}{k_i \prod_{j=1}^{i-1} K_j}\right) J = [S_{in}] - \frac{[S_{out}]}{\prod_{i=1}^{n} K_i} \tag{9}$$

Thus the steady state pathway flux $J$ can be computed as:

$$J = \frac{[S_{in}] - \frac{[S_{out}]}{\prod_{i=1}^{n} K_i}}{\sum_{i=1}^{n} \frac{1}{k_i \prod_{j=1}^{i-1} K_j}} \tag{10}$$

##### 1.3.2 Zero-order kinetics

It is worth noting that under zero-order kinetics, we cannot derive an analytical expression to calculate the steady state pathway flux from the kinetic and thermodynamic parameters and concentrations of input and output metabolites. This is because the steady state equations converge to a high degree polynomial equation of $J$, which does not have an algebraic solution when the pathway contains more than five reactions. Hence, here we only derive the steady state equation with respect to the pathway flux $J$ without providing analytical expression for its solution.

First, we write the steady-state equations of the linear metabolic pathway with zero-order kinetics:

$$\begin{cases} J = k_1 \left(1 - \frac{[S_1]}{K_1 [S_{in}]}\right) \\ J = k_2 \left(1 - \frac{[S_2]}{K_2 [S_1]}\right) \\ \vdots \\ J = k_i \left(1 - \frac{[S_i]}{K_i [S_{i-1}]}\right) \\ \vdots \\ J = k_n \left(1 - \frac{[S_{out}]}{K_n [S_{n-1}]}\right) \end{cases} \tag{11}$$

The equations can be reshaped as below:

$$\begin{cases} \frac{[S_1]}{[S_{in}]} = K_1 \left(1 - \frac{J}{k_1}\right) \\ \frac{[S_2]}{[S_1]} = K_2 \left(1 - \frac{J}{k_2}\right) \\ \vdots \\ \frac{[S_i]}{[S_{i-1}]} = K_i \left(1 - \frac{J}{k_i}\right) \\ \vdots \\ \frac{[S_{out}]}{[S_{n-1}]} = K_n \left(1 - \frac{J}{k_n}\right) \end{cases} \tag{12}$$

Multiplying both sides of all equations, we have:

$$\frac{[S_{out}]}{[S_{in}]} = \prod_{i=1}^{n} K_i(1 - \frac{J}{k_i}) \tag{13}$$

This is an algebraic equation of $J$ with degree $n$, in which $n$ is the number of reactions in the pathway. From the Abel-Ruffini theorem, this kind of equation does not have an algebraic solution (i.e. an analytical expression of $J$ as a function of the parameters $\mathbf{k}, \mathbf{K}, [S_{in}]$ and $[S_{out}]$).

#### 1.4 Flux control efficients

Flux control coefficients (FCCs) quantify how the pathway flux $J$ responds to changes in the enzyme catalyzing a single reaction in the pathway. As we have previously derived[1], under first-order kinetics, the scaled flux control coefficient (FCC) for the $i$-th reaction in the linear pathway can be computed from the kinetic and thermodynamic parameters using the equation below:

$$C^J_{v_i} = \frac{\partial \log J}{\partial \log k_i} = \frac{1}{k_i \prod_{l=1}^{i-1} K_l (\sum_{l=1}^{n} \frac{1}{k_l \prod_{m=1}^{l-1} K_m})} \tag{14}$$

In other words, we have:

$$C^J_{v_i} \propto \frac{1}{k_i \prod_{l=1}^{i-1} K_l} \tag{15}$$

According to this equation, we can also compute the ratio between two FCCs $C^J_{v_i}$ and $C^J_{v_{i+1}}$:

$$\frac{C^J_{v_i}}{C^J_{v_{i+1}}} = \frac{k_{i+1} K_i}{k_i} \tag{16}$$

For the zero-order kinetics, as the zero-order steady state equations are high-degree polynomial equations, it is difficult to derive analytical expressions that compute the FCCs as explicit functions of the kinetic and thermodynamic parameters. However, Noor et al. have derived that the FCCs can be written as explicit functions of the scaled Gibbs free energy change $g$ of each reaction:

$$C^J_{v_i} = \frac{e^{-g_i} - 1}{\sum_{j=1}^{n} (e^{-g_j} - 1)} \tag{17}$$

That relationship can be derived by applying the summation and connectivity theorems in the theory of metabolic control analysis. Briefly, the summation theorem guarantees the sum of all FCCs in the pathway equals one:

$$\sum_{i=1}^{n} C^J_{v_i} = 1 \tag{18}$$

And the connectivity theorem guarantees that the pathway flux is invariant upon perturbation to concentrations of intermediate metabolites (i.e. metabolites that are neither $S_{in}$ nor $S_{out}$). The mathematical expressions of the connectivity theorem depend on the definition of metabolite elasticity coefficients, which quantify the local responses of reaction velocities to changes in the concentrations of substrate or product. The term "local" means that in computing the elasticity coefficients, reactions that do not directly consume or produce the metabolite are not taken into consideration. In the linear metabolic pathway, the concentration of the $i$-th metabolite is locally connected to the velocities of the $i$-th and $i+1$-th reactions, thereby we have the two corresponding elasticity coefficients:

$$\begin{cases} \epsilon_{S_i}^{v_i} = \dfrac{\partial \log v_i}{\partial \log [S_i]} = -\dfrac{S_i}{K_i v_i [S_{i-1}]} \\ \epsilon_{S_i}^{v_{i+1}} = \dfrac{\partial \log v_{i+1}}{\partial \log [S_i]} = \dfrac{k_{i+1}[S_{i+1}]}{K_{i+1} v_{i+1} [S_i]} \end{cases} \tag{19}$$

According to the connectivity theorem:

$$C_{v_i}^J \epsilon_{S_i}^{v_i} + C_{v_{i+1}}^J \epsilon_{S_i}^{v_{i+1}} = 0 \tag{20}$$

The ratio between any two subsequent FCCs in the linear metabolic pathway can be then written as:

$$\frac{C_{v_{i+1}}^J}{C_{v_i}^J} = \frac{k_i K_{i+1} [S_i]^2}{K_i k_{i+1} [S_{i-1}][S_{i+1}]} \tag{21}$$

This expression can be simplified by introducing terms for reaction thermodynamics. According to (4), we can define thermodynamics-related terms below:

$$g_i = \log \frac{[S_i]}{K_i [S_{i-1}]} \tag{22}$$

Combining (21) and (22), we have:

$$\frac{C_{v_{i+1}}^J}{C_{v_i}^J} = \frac{k_i e^{g_i}}{k_{i+1} e^{g_{i+1}}} \tag{23}$$

Furthermore, at the steady state, the velocities of the $i$-th and $i+1$-th reactions should be the same, therefore we have the following equation that links the apparent rate constants and scaled reaction Gibbs free energy changes of different reactions:

$$k_i(1 - e^{g_i}) = k_{i+1}(1 - e^{g_{i+1}}) \tag{24}$$

Therefore, the ratio of $C_{v_{i+1}}^J$ to $C_{v_i}^J$ can be determined by the scaled reaction Gibbs free energy changes:

$$\frac{C_{v_{i+1}}^J}{C_{v_i}^J} = \frac{e^{-g_{i+1}} - 1}{e^{-g_i} - 1} \tag{25}$$

Finally, as the sum of all FCCs in the pathway equals to one, each FCC can be computed as the function of the scaled Gibbs free energy changes:

$$C_{v_i}^J = \frac{e^{-g_i} - 1}{\sum_{j=1}^{n} (e^{-g_j} - 1)} \tag{26}$$

### 2 Maximization of flux efficiency in linear metabolic pathways

#### 2.1 Definition and maximization of flux efficiency

To define the flux efficiency of the metabolic pathway, we first introduce relationship between enzyme abundance and apparent rate constant in our model:

$$k_i = a_i e_i \tag{27}$$

In which $k_i$ is the apparent rate constant as described previously, $e_i$ is the abundance of enzyme catalyzing this reaction, $a_i$ is a linear coefficient that can be interpreted as the catalytic efficiency of that enzyme. The enzyme cost of the pathway is then defined as:

$$E = \sum_{i=1}^{n} e_i \tag{28}$$

With the definition of the catalytic efficiencies a $= \{a_i\}$ and enzyme abundances e $= \{e_i\}$, the pathway flux $J$ can be also written as a function of these terms:

$$J = J(\mathbf{a}, \mathbf{e}, \mathbf{K}, S_{in}, S_{out}) \tag{29}$$

We therefore define the flux efficiency of a linear metabolic pathway as the ratio of the pathway flux $J$ to its enzyme cost $E$:

$$\eta = \frac{J(\mathbf{a}, \mathbf{e}, \mathbf{K}, S_{in}, S_{out})}{\sum_{i=1}^{n} e_i} \tag{30}$$

Since the pathway flux $J$ is linear to the apparent rate constant $\mathbf{k}$, which is linear to the enzyme abundances $e$, the flux efficiency $\eta$ is invariant upon scaling $\mathbf{e}$:

$$\eta(\mathbf{a}, \mathbf{e}, \mathbf{K}, S_{in}, S_{out}) = \eta(\mathbf{a}, \alpha\mathbf{e}, \mathbf{K}, S_{in}, S_{out}) \tag{31}$$

Hence, the problem of maximizing flux efficiency is equivalent to the problem of minimizing enzyme cost, which can be written as below:

$$\begin{gathered} \min \sum_{i=1}^{n} e_i, \text{ s.t.} \\ J_{\mathbf{a},\mathbf{K},S_{in},S_{out}}(\mathbf{e}) = J_0, \ e_i \geq 0 \end{gathered} \tag{32}$$

To eliminate the inequality constraints, let $e_i = x_i^2$, in other words, $\mathbf{e} = \mathbf{x} \circ \mathbf{x}$. The constrained optimization is thus converted to the form below:

$$\begin{gathered} \min \sum_{i=1}^{n} x_i^2, \text{ s.t.} \\ J_{\mathbf{a},\mathbf{K},S_{in},S_{out}}(\mathbf{x} \circ \mathbf{x}) = J_0 \end{gathered} \tag{33}$$

Next, define the Lagrange function as below:

$$L(\mathbf{x}, \lambda) = \sum_{i=1}^{n} x_i^2 - \lambda(J_{\mathbf{a},\mathbf{K},S_{in},S_{out}}(\mathbf{x} \circ \mathbf{x}) - J_0) \tag{34}$$

The optimal solution $\mathbf{x}_{opt}$ that minimizes the enzyme cost thus satisfies the conditions below:

$$\begin{cases} \dfrac{\partial L}{\partial x_i} = 0 \\ \dfrac{\partial L}{\partial \lambda} = 0 \end{cases} \tag{35}$$

In which the partial derivatives with respect to $x_i$ can be computed:

$$\frac{\partial L}{\partial x_i} = 2x_i \left(1 - \lambda a_i \frac{\partial J}{\partial k_i}\right) \tag{36}$$

Therefore $\frac{\partial J}{\partial k_i} \propto \frac{1}{a_i}$ is required for the solution minimizing the enzyme cost. Hence we have:

$$C^J_{v_i} = \frac{k_i}{J}\frac{\partial J}{\partial k_i} \propto \frac{k_i}{a_i} \tag{37}$$

As $k_i = a_i e_i$, we have:

$$C^J_{v_i} \propto e_i \tag{38}$$

Which means that when the total enzyme cost is minimized, the FCCs are linear to the abundance of enzymes catalyzing the corresponding reactions. This is consistent with the enzyme-control rule reported in previous studies.

##### 2.1.1 First-order kinetics

For linear pathways with first-order kinetics, we can combine (16) and (37) to derive analytical relationships between the optimal solution $\mathbf{e}$ and other parameters in the model. First, according to (37):

$$\frac{C^J_{v_i}}{C^J_{v_{i+1}}} = \frac{e_i}{e_{i+1}} \tag{39}$$

Combining that relationship with (16), we have:

$$\frac{K_i k_{i+1}}{k_i} = \frac{e_i}{e_{i+1}} \tag{40}$$

Therefore, the catalytic efficiencies, enzyme abundances, and equilibrium constants are connected by the relationship below:

$$K_i = \frac{a_i e_i^2}{a_{i+1} e_{i+1}^2} \tag{41}$$

##### 2.1.2 Zero-order kinetics

For zero-order kinetics, for any two reactions in the pathway we have the three equations below:

$$\begin{cases} \dfrac{C^J_{v_i}}{C^J_{v_j}} = \dfrac{e^{-g_i} - 1}{e^{-g_j} - 1} \\ \dfrac{C^J_{v_i}}{C^J_{v_j}} = \dfrac{e_i}{e_j} \\ a_i e_i (1 - e^{g_i}) = a_j e_j (1 - e^{g_j}) \end{cases} \tag{42}$$

The first equation reflects the relationship between reaction thermodynamics and FCCs in linear metabolic pathways with zero-order kinetics that we have derived in (26), the second equation is the enzyme-control rule when the enzyme cost is minimized, and the last equation means that any two reaction velocities equal with each other in the steady state. Combining the three reactions, we have:

$$e^{-(g_i - g_j)} = \frac{a_i e_i^2}{a_j e_j^2}, \forall i, j \tag{43}$$

If we let $j = i + 1$, because $e^{-g_i} = \frac{K_i[S_{i-1}]}{[S_i]}$, this relationship will have similar form to that for the first-order kinetics, in which the ratio between $a_i e_i^2$ and $a_{i+1} e_{i+1}^2$ is determined by a thermodynamics-related term:

$$K_i \frac{[S_{i-1}][S_{i+1}]}{K_{i+1}[S_i]^2} = \frac{a_i e_i^2}{a_{i+1} e_{i+1}^2} \tag{44}$$

In the case of first-order kinetics, the thermodynamic term only depends on the equilibrium constant or standard Gibbs free energy change of a reaction, while in the case of zero-order kinetics, it relies on the reaction Gibbs free energy that is determined by the concentrations of metabolites as well as the standard Gibbs free energy change. Let $\Phi_i = \frac{[S_{i-1}][S_{i+1}]}{K_{i+1}[S_i]^2}$, the relationship has the form below:

$$K_i \Phi_i = \frac{a_i e_i^2}{a_{i+1} e_{i+1}^2} \tag{45}$$

##### 2.1.3 Treatment of isozymes

In real metabolic networks, it is a frequent case where one reaction is catalyzed by several isozymes. In this case, the relationship between the apparent rate constant $\mathbf{k}$ and enzyme abundance $\mathbf{e}$ becomes:

$$k_i = \sum_{j=1}^{n_i} a_{ij} e_{ij} \tag{46}$$

In which $n_i$ is the number of isozymes catalyzing the $i$-th reaction, $a_{ij}$ is the catalytic efficiency of the $j$-th isozyme for the $i$-th reaction, $e_{ij}$ is the abundance of this isozyme. The total enzyme cost of a pathway then becomes:

$$E = \sum_{i=1}^{n} \sum_{j=1}^{n_i} e_{ij} \tag{47}$$

It is worth noting that if we allow each $e_{ij}$ to be freely optimized in maximization of the flux efficiency, for a given reaction $i$, only the isozyme with largest $a_{ij}$ will have non-zero abundance under optimal flux efficiency, because replacing any other isozyme with this max-efficiency isozyme of idential abundance will increase the flux efficiency $\eta$. Therefore, we assume that the fractions of the isozymes $e_{ij}$ in the total abundance of enzymes catalyzing reaction $i$ are fixed, that is:

$$e_{ij} = c_{ij} e_i \tag{48}$$

In which $c_{ij}$ is the fraction of the $j$-th isozyme in all isozymes catalyzing the $i$-th reaction. Therefore, we have the relationship below:

$$k_i = e_i \sum_{j=1}^{n_i} a_{ij} c_{ij} \tag{49}$$

This relationship gives a definition of "apparent catalytic efficiency" of enzymes catalyzing the reaction $i$, which is a weighted average of the catalytic efficiencies of its isozymes:

$$\hat{a}_i = \sum_{j=1}^{n_i} a_{ij} c_{ij} \tag{50}$$

Therefore, the optimal solution for maximizing flux efficiency under the first-order kinetics satisfies the relationship below:

$$K_i = \frac{\sum_{j=1}^{n_i} a_{ij} c_{ij} e_i^2}{\sum_{j=1}^{n_{i+1}} a_{i+1,j} c_{i+1,j} e_{i+1}^2} \tag{51}$$

Substituting $e_i = \sum_{i=1}^{n_i} e_{ij}$ into this equation, we have the following relationship:

$$K_i = \frac{\sum_{j=1}^{n_i} e_{ij} \sum_{j=1}^{n_i} a_{ij} e_{ij}}{\sum_{j=1}^{n_{i+1}} e_{i+1,j} \sum_{j=1}^{n_{i+1}} a_{i+1,j} e_{i+1,j}} \tag{52}$$

For zero-order kinetics, this relationship becomes:

$$e^{-(g_i - g_j)} = \frac{\sum_{k=1}^{n_i} e_{ik} \sum_{k=1}^{n_i} a_{ik} e_{ik}}{\sum_{k=1}^{n_j} e_{jk} \sum_{k=1}^{n_j} a_{jk} e_{jk}}, \forall i, j \tag{53}$$

#### 2.2 Conditions for optimal flux efficiency under Michaelis-Menten kinetics

Here we consider the reversible Michaelis-Menten reaction kinetics for a unimolecular metabolic reaction:

$$v = \frac{k_{cat}[E]\frac{[S]}{K_S}}{1 + \frac{[S]}{K_S} + \frac{[P]}{K_P}} \left(1 - \frac{[P]}{K[S]}\right) \tag{54}$$

In which $k_{cat}$ is the turnover number of the enzyme, $K_S$ and $K_P$ are the Michaelis constants for the substrate $S$ and product $P$, $K$ is the equilibrium constant of the reaction.

We have proved previously that under optimal flux efficiency, the flux control coefficients are proportional to enzyme abundances: $C_{v_i}^J \propto e_i$. This condition is independent of the kinetic rule of the reaction[2]. Therefore, local relationships linking the enzyme abundances, metabolite concentrations, kinetic and thermodynamic constants can be derived using this relationship and the connectivity theorem under the framework of metabolic control analysis.

For the Michaelis-Menten kinetics, we first derive analytical expressions of its elasticity coefficients:

$$\begin{cases} \epsilon_S = \dfrac{\partial \log v}{\partial \log [S]} & = -\dfrac{[S](K_P[P] + K_S K[S]([P] + K_P))}{([P] - K[S])(K_S[P] + K_P(K_S + [S]))} \\ \epsilon_P = \dfrac{\partial \log v}{\partial \log [P]} & = \dfrac{[P](K_S K[S] + K_P(K_S + [S]))}{([P] - K[S])(K_S[P] + K_P(K_S + [S]))} \end{cases} \tag{55}$$

When the Michaelis constants $K_S$ and $K_P$ are much higher than the concentrations $[S]$ and $[P]$, the elasticity coefficients converge to those under first-order kinetics:

$$\begin{cases} \lim\limits_{K_S, K_P \to +\infty} \epsilon_S = -\dfrac{K[S]}{[P] - K[S]} \\ \lim\limits_{K_S, K_P \to +\infty} \epsilon_P = \dfrac{[P]}{[P] - K[S]} \end{cases} \tag{56}$$

Let $\epsilon_S^{(1)}$ and $\epsilon_P^{(1)}$ denote the elasticity coefficients under first-order kinetics, we can therefore decouple the elasticity coefficients for Michaelis-Menten kinetics into the product of two terms:

$$\begin{cases} \epsilon_S = \epsilon_S^{(1)} \Psi_S \\ \epsilon_P = \epsilon_P^{(1)} \Psi_P \end{cases} \tag{57}$$

In which:

$$\begin{cases} \Psi_S = \dfrac{K_P[P] + K_S K[S]([P] + K_P)}{K_S K[P] + K_P K(K_S + [S])} \\ \Psi_P = \dfrac{K_S K[S] + K_P(K_S + [S])}{K_S[P] + K_P(K_S + [S])} \\ \epsilon_S^{(1)} = -\dfrac{K[S]}{[P] - K[S]} \\ \epsilon_P^{(1)} = \dfrac{[P]}{[P] - K[S]} \end{cases} \tag{58}$$

Note that $\epsilon_S^{(1)}$ and $\epsilon_P^{(1)}$ are the elasticity coefficients for 1st-order kinetics. This relationship also holds for a linear reaction chain:

$$\begin{cases} \epsilon_{S_i}^{v_{i+1}} = \epsilon_{S_i}^{v_{i+1}(1)} \Psi_{S_i}^{v_{i+1}} \\ \epsilon_{S_{i+1}}^{v_{i+1}} = \epsilon_{S_{i+1}}^{v_{i+1}(1)} \Psi_{S_{i+1}}^{v_{i+1}} \end{cases} \tag{59}$$

Furthermore, according to the connectivity theorem:

$$C_{v_i}^J \epsilon_{S_i}^{v_i} + C_{v_{i+1}}^J \epsilon_{S_i}^{v_{i+1}} = 0 \tag{60}$$

As the relationship $C_{v_i}^J \propto e_i$ holds when the pathway achieves optimal flux efficiency, we have the relationship below:

$$e_i \epsilon_{S_i}^{v_i} + e_{i+1} \epsilon_{S_i}^{v_{i+1}} = 0 \tag{61}$$

Therefore, the ratio of the enzyme abundances $e_i$ to $e_{i+1}$ can be computed as follows:

$$\frac{e_i}{e_{i+1}} = -\frac{\epsilon_{S_i}^{v_{i+1}}}{\epsilon_{S_i}^{v_i}} = -\frac{\epsilon_{S_i}^{v_{i+1}(1)} \Psi_{S_i}^{v_{i+1}}}{\epsilon_{S_i}^{v_i(1)} \Psi_{S_i}^{v_i}} \tag{62}$$

Let $\Phi_i = \frac{\Psi_{S_i}^{v_{i+1}}}{\Psi_{S_i}^{v_i}}$, then subject the expressions of $\epsilon_{S_i}^{v_i(1)}$ and $\epsilon_{S_i}^{v_{i+1}(1)}$ into the equation above, we have:

$$\frac{e_i}{e_{i+1}} = K_{i+1} \Phi_i \frac{[S_i] - K_i[S_{i-1}]}{[S_{i+1}] - K_{i+1}[S_i]} \tag{63}$$

Let $a = \frac{k_{cat}/K_S}{1+[S]/K_S+[P]/K_P}$, the reaction rate under Michaelis-Menten kinetics can be written as:

$$v = a[E]([S] - \frac{[P]}{K}) \tag{64}$$

For reactions in the linear metabolic pathway, reaction rates can also be written in this form:

$$v_i = a_i e_i \left( [S_{i-1}] - \frac{[S_i]}{K_i} \right) \tag{65}$$

In which $a_i = \frac{k_{cat,i}/K_{S,i}}{1+[S_{i-1}]/K_{S,i}+[S_i]/K_{P,i}}$. Since the metabolic pathway is at steady state, we have $v_i = v_{i+1}$. Therefore:

$$a_i e_i \left( [S_{i-1}] - \frac{[S_i]}{K_i} \right) = a_{i+1} e_{i+1} \left( [S_i] - \frac{[S_{i+1}]}{K_{i+1}} \right) \tag{66}$$

Hence the ratio of $e_i$ to $e_{i+1}$ can be written as follows:

$$\frac{e_i}{e_{i+1}} = \frac{K_i a_{i+1}([S_{i+1}] - K_{i+1}[S_i])}{K_{i+1} a_i([S_i] - K_i[S_{i-1}])} \tag{67}$$

Finally, combining (63) and (67), we have the relationship:

$$K_i \Phi_i = \frac{a_i e_i^2}{a_{i+1} e_{i+1}^2} \tag{68}$$

### 3 Optimization of efficiency in branching pathways

#### 3.1 Scheme of branching pathways

Here we consider two simplest topologies of branching metabolic pathways. The first is a diverging pathway with one upstream reaction and $n-1$ downstream reactions, connected by a branching point metabolite $S_b$:

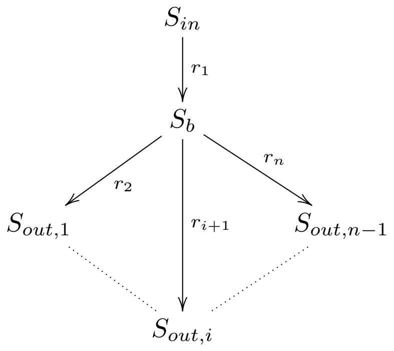


(69)

The other one is a converging pathway with $n-1$ upstream reactions and one downstream reactions, connected by a branching point metabolite $S_b$.

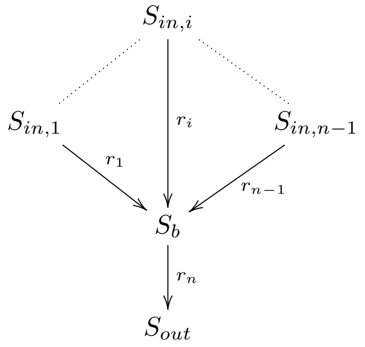


(70)

Similar to the case of linear pathways, each reaction in the branching pathways also has an apparent rate constant $k$ and an equilibrium constant $K$. The kinetic and thermodynamic parameters for the $i$-th reaction, $r_i$, is referred to as $\{k_i, K_i\}$. For metabolites included in these branching pathways, only the metabolite at the branching point has its concentration as a free variable, while concentrations of all upstream and downstream metabolites, $[S_{in,i}]$ and $[S_{out,i}]$, are treated as boundary conditions. For a given pathway, the equilibrium constants $\mathbf{K}$ are determined by standard Gibbs free energy changes of the reactions, thereby these are also treated as constants.

#### 3.2 Relationship between k and $[S_b]$

For the branching pathways, it is worth noting that when the steady state flux configuration $\mathbf{J} = \{J_1, \cdots, J_n\}$ is known, each apparent rate constant $k_i$ can be written as a function of the parameters $[\mathbf{S}_{in}]$, $[\mathbf{S}_{out}]$, $\mathbf{K}$, and the concentration of the branching point metabolite, $[S_b]$. For pathways with first-order kinetics, these expressions are provided below:

##### 3.2.1 Diverging pathway

For the diverging pathway, $J_1$ is the flux carried by the upstream reaction, and $J_2$ to $J_n$ are fluxes of the downstream reactions. At the steady state we have $J_1 = \sum_{i=2}^{n} J_i$. Therefore, we have the equations below:

$$\begin{cases} k_1 = \dfrac{K_1 \sum_{i=2}^{n} J_i}{K_1[S_{in}] - [S_b]} \\ k_2 = \dfrac{K_2 J_2}{K_2[S_b] - [S_{out,1}]} \\ \quad\vdots \\ k_i = \dfrac{K_i J_i}{K_i[S_b] - [S_{out,i-1}]} \\ \quad\vdots \\ k_n = \dfrac{K_n J_n}{K_n[S_b] - [S_{out,n-1}]} \end{cases} \tag{71}$$

##### 3.2.2 Converging pathway

For the converging pathway, $J_1$ to $J_{n-1}$ are fluxes of the upstream reactions that converge to one downstream flux $J_n$. At the steady state, we have $J_n = \sum_{i=1}^{n-1} J_i$, which allows us to derive the equations below:

$$\begin{cases} k_1 = \dfrac{K_1 J_1}{K_1[S_{in,1}] - [S_b]} \\ \quad\vdots \\ k_i = \dfrac{K_i J_i}{K_i[S_{in,i}] - [S_b]} \\ \quad\vdots \\ k_{n-1} = \dfrac{K_{n-1} J_{n-1}}{K_{n-1}[S_{in,n-1}] - [S_b]} \\ k_n = \dfrac{K_n \sum_{i=1}^{n-1} J_i}{K_n[S_b] - [S_{out}]} \end{cases} \tag{72}$$

#### 3.3 Minimization of enzyme cost

Here, we consider the problem of minimizing enzyme cost in branching pathways. The form of the optimization problem is quite similar to that of minimizing $\sum_{i=1}^{n} k_i$. Assuming that for each reaction

$k_i = a_i e_i$, in which $e_i$ is the abundance of enzyme catalyzing the $i$-reaction, the problem becomes the minimization of $\sum_{i=1}^{n} e_i = \sum_{i=1}^{n} \frac{k_i}{a_i}$:

$$\min e = \sum_{i=1}^{n} \frac{k_i}{a_i}, \text{ s.t. } \mathbf{J} = \mathbf{J}_0 \tag{73}$$

##### 3.3.1 Diverging pathway

For the diverging pathway, the objective function $\sum_{i=1}^{n} \frac{k_i}{a_i}$ can be written as a function of the intermediate metabolite concentration $[S_b]$:

$$\sum_{i=1}^{n} e_i = \sum_{i=1}^{n} \frac{k_i}{a_i} = \frac{K_1 \sum_{i=2}^{n} J_i}{a_1 \left(K_1[S_{in}] - [S_b]\right)} + \sum_{i=2}^{n} \frac{K_i J_i}{a_i \left(K_i[S_b] - [S_{out,i-1}]\right)} \tag{74}$$

We compute the derivative of this objective function with respect to $[S_b]$ to identify potential optimal solutions:

$$\frac{\partial(\sum_{i=1}^{n} e_i)}{\partial[S_b]} = \frac{K_1 \sum_{i=2}^{n} J_i}{a_1(K_1[S_{in}] - [S_b])^2} - \sum_{i=2}^{n} \frac{K_i^2 J_i}{a_i(K_i[S_b] - [S_{out,i-1}])^2} \tag{75}$$

At the critical point, the derivative equals zero, therefore we have:

$$\frac{K_1 \sum_{i=2}^{n} J_i}{a_1(K_1[S_{in}] - [S_b])^2} = \sum_{i=2}^{n} \frac{K_i^2 J_i}{a_i(K_i[S_b] - [S_{out,i-1}])^2} \tag{76}$$

Substituting the expressions of $k_i$'s as functions of $[S_b]$ to elinimate the concentration terms, we have:

$$\frac{k_1^2}{a_1 \left(K_1 \sum_{i=2}^{n} J_i\right)} = \sum_{i=2}^{n} \frac{k_i^2}{a_i J_i} \tag{77}$$

Since $k_i = a_i e_i$, we have:

$$\frac{a_1 e_1^2}{K_1 \sum_{i=2}^{n} J_i} = \sum_{i=2}^{n} \frac{a_i e_i^2}{J_i} \tag{78}$$

Hence we have:

$$\frac{a_1 e_1^2}{K_1} = \left(\sum_{i=2}^{n} J_i\right) \left(\sum_{i=2}^{n} \frac{a_i e_i^2}{J_i}\right) > \sum_{i=2}^{n} a_i e_i^2 \tag{79}$$

Therefore the inequality below holds for any flux configuration $\mathbf{J}$:

$$K_1 < \frac{a_1 e_1^2}{\sum_{i=2}^{n} a_i e_i^2} \tag{80}$$

##### 3.3.2 Converging pathway

For the converging pathway, we also compute the derivative of $\sum_{i=1}^{n} e_i$ with respect to $[S_b]$ to identify potential optimal solutions:

$$\frac{\partial(\sum_{i=1}^{n} e_i)}{\partial[S_b]} = \sum_{i=1}^{n-1} \frac{K_i J_i}{a_i(K_i[S_{in,i}] - [S_b])^2} - \frac{K_n^2 \sum_{i=1}^{n-1} J_i}{a_n(K_n[S_b] - [S_{out}])^2} \tag{81}$$

At the critical point the derivative equals zero, therefore we have:

$$\sum_{i=1}^{n-1} \frac{K_i J_i}{a_i(K_i[S_{in,i}] - [S_b])^2} = \frac{K_n^2 \sum_{i=1}^{n-1} J_i}{a_n(K_n[S_b] - [S_{out}])^2} \tag{82}$$

Again, we substitute the expressions of $k_i$'s as functions of $[S_b]$ to elinimate the concentration terms:

$$\sum_{i=1}^{n-1} \frac{k_i^2}{a_i K_i J_i} = \frac{k_n^2}{a_n \sum_{i=1}^{n-1} J_i} \tag{83}$$

Since $k_i = a_i e_i$, we have:

$$\sum_{i=1}^{n-1} \frac{a_i e_i^2}{K_i J_i} = \frac{a_n e_n^2}{\sum_{i=1}^{n-1} J_i} \tag{84}$$

Therefore, we have:

$$a_n e_n^2 = \left( \sum_{i=1}^{n-1} \frac{a_i e_i^2}{K_i J_i} \right) \left( \sum_{i=1}^{n-1} J_i \right) > \sum_{i=1}^{n-1} \frac{a_i e_i^2}{K_i} \tag{85}$$

Hence the inequality below holds for any flux configuration $\mathbf{J}$:

$$\sum_{i=1}^{n-1} \frac{a_i e_i^2}{K_i} < a_n e_n^2 \tag{86}$$

### References


[1] Z. Dai and J. W. Locasale. Thermodynamic constraints on the regulation of metabolic fluxes. *J Biol Chem*, 293(51):19725–19739, 2018. Dai, Ziwei Locasale, Jason W eng R00 CA168997/CA/NCI NIH HHS/ R01 CA193256/CA/NCI NIH HHS/ Research Support, N.I.H., Extramural Research Support, Non-U.S. Gov't 2018/10/27 J Biol Chem. 2018 Dec 21;293(51):19725-19739. doi: 10.1074/jbc.RA118.004372. Epub 2018 Oct 25.

[2] Reinhart Heinrich and Edda Klipp. Control Analysis of Unbranched Enzymatic Chains in States of Maximal Activity. *Journal of Theoretical Biology*, 182(3):243–252, October 1996.
