## Supplementary Figures for "A universal law of thermodynamics-kinetics coupling shapes enzyme allocation and glycolytic efficiency across species"

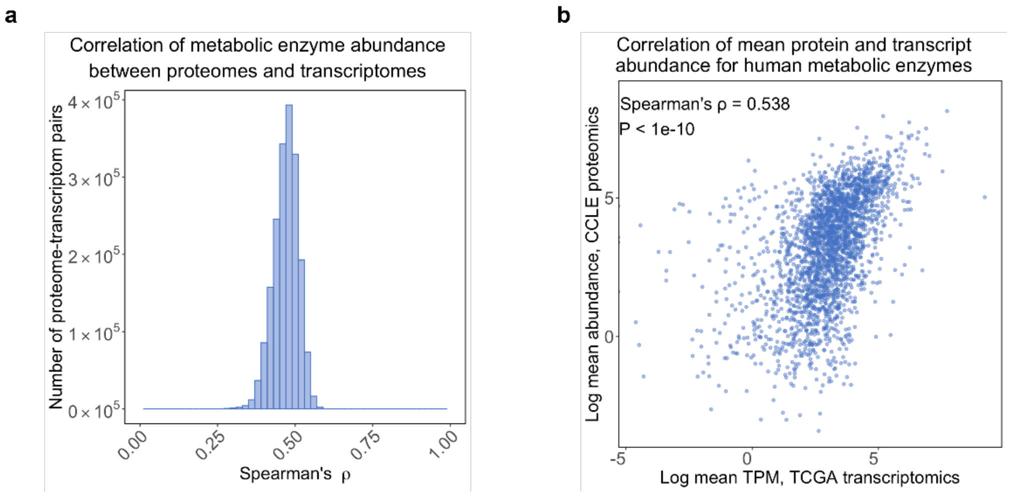


**Figure S1 (Related to Figure 1). Correlation between protein and transcript abundances of human metabolic enzymes.** (a) Distribution of Spearman correlation between protein abundance in CCLE cell lines and transcript abundance in TCGA tumors. (b) Scatter plot comparing mean TPM of the TCGA transcriptomics data and mean protein abundance in the CCLE proteomics data.

**a**

| | First-order | Zero-order | Michaelis-Menten |
|---|---|---|---|
| **Reaction kinetics** | $v = a[E]\left([S] - \frac{[P]}{K}\right)$ | $v = a[E]\left(1 - \frac{[P]}{K[S]}\right)$ | $v = k_{cat}[E]\frac{\frac{[S]}{K_s}}{1+\frac{[S]}{K_s}+\frac{[P]}{K_p}}\left(1 - \frac{[P]}{K[S]}\right)$ |
| **Condition for optimal flux efficiency** | $K_i = \frac{a_i[E_i]^2}{a_{i+1}[E_{i+1}]^2}$ | $e^{-(g_i - g_j)} = \frac{a_i[E_i]^2}{a_j[E_j]^2}$ | $K_i\Phi_i = \frac{a_i[E_i]^2}{a_{i+1}[E_{i+1}]^2}$ |
| **Condition for optimal flux efficiency, generalized form** | $\Phi_i = 1$ | $\Phi_i = \frac{[S_{i-1}][S_{i+1}]}{K_{i+1}[S_i]^2}$ | $\Phi_i = \frac{\Psi_{S_i}^{v_{i+1}}}{\Psi_{S_i}^{v_i}}$ <br> $\Psi_{S_i}^{v_{i+1}} = \frac{K_{P,i+1}[S_{i+1}] + K_{S,i+1}K_{i+1}[S_i]([S_{i+1}] + K_{P,i+1})}{K_{S,i+1}K_{i+1}[S_{i+1}] + K_{P,i+1}K_{i+1}(K_{S,i+1} + [S_i])}$ <br> $\Psi_{S_i}^{v_i} = \frac{K_{S,i}K_i[S_{i-1}] + K_{P,i}(K_{S,i} + [S_{i-1}])}{K_{S,i}[S_i] + K_{P,i}(K_{S,i} + [S_{i-1}])}$ |

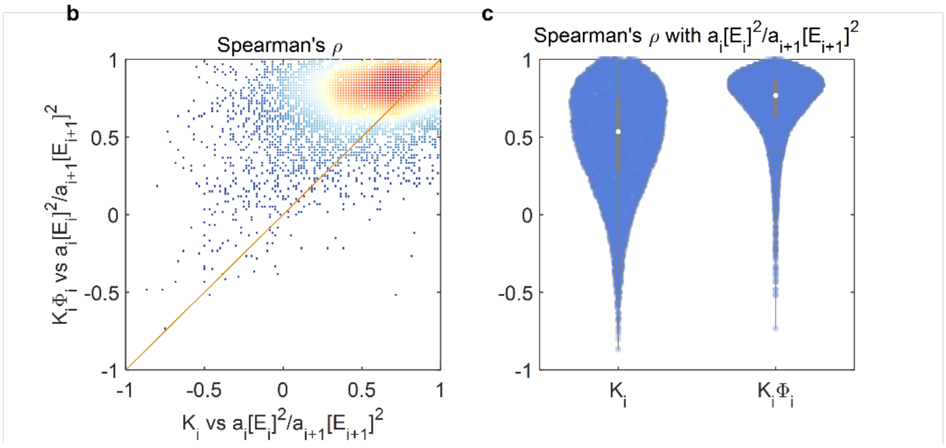


**Figure S2 (Related to Figure 2). Optimal conditions for maximal flux efficiency under different rate laws.** (a) Conditions for optimal flux efficiency under first-order, zero-order, and Michaelis-Menten kinetics. The generalized form for the optimal condition is $K_i\Phi_i = \frac{a_i[E_i]^2}{a_{i+1}[E_{i+1}]^2}$. (b) Density scatter plot comparing the Spearman correlation of $K_i$ or $K_i\Phi_i$ with the CAQ, $\frac{a_i[E_i]^2}{a_{i+1}[E_{i+1}]^2}$, for optimal enzyme abundances maximizing the flux efficiency of a 10-step linear pathway with Michaelis-Menten kinetics. (c) Violin plot comparing the Spearman correlation of $K_i$ or $K_i\Phi_i$ with the CAQ, $\frac{a_i[E_i]^2}{a_{i+1}[E_{i+1}]^2}$, for optimal enzyme abundances maximizing the flux efficiency of a 10-step linear pathway with Michaelis-Menten kinetics.

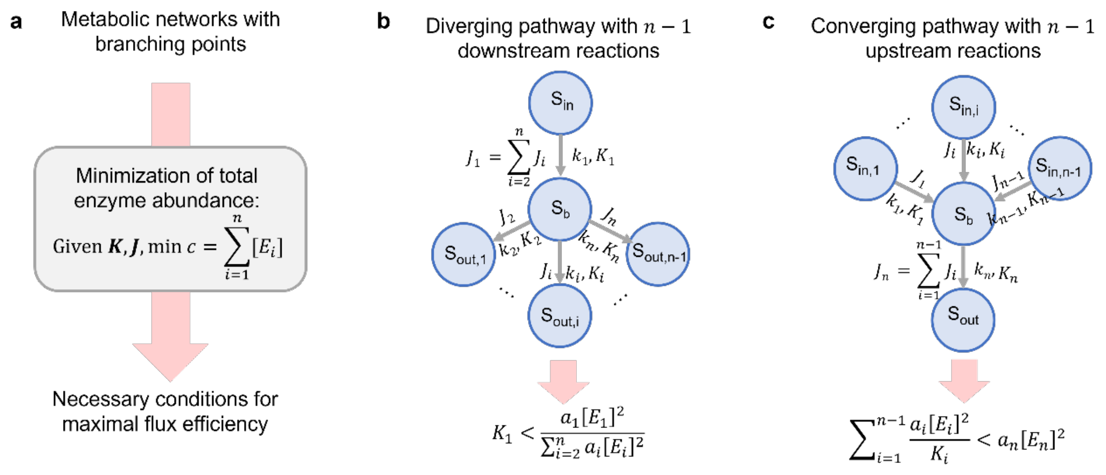


**Figure S3 (Related to Figure 2). Conditions for optimal flux efficiency in branching pathways.** (a) Mathematical definition of the optimization problem. (b) Scheme of the diverging pathway and the corresponding condition for optimal efficiency. (c) Scheme of the converging pathway and the corresponding condition for optimal efficiency.

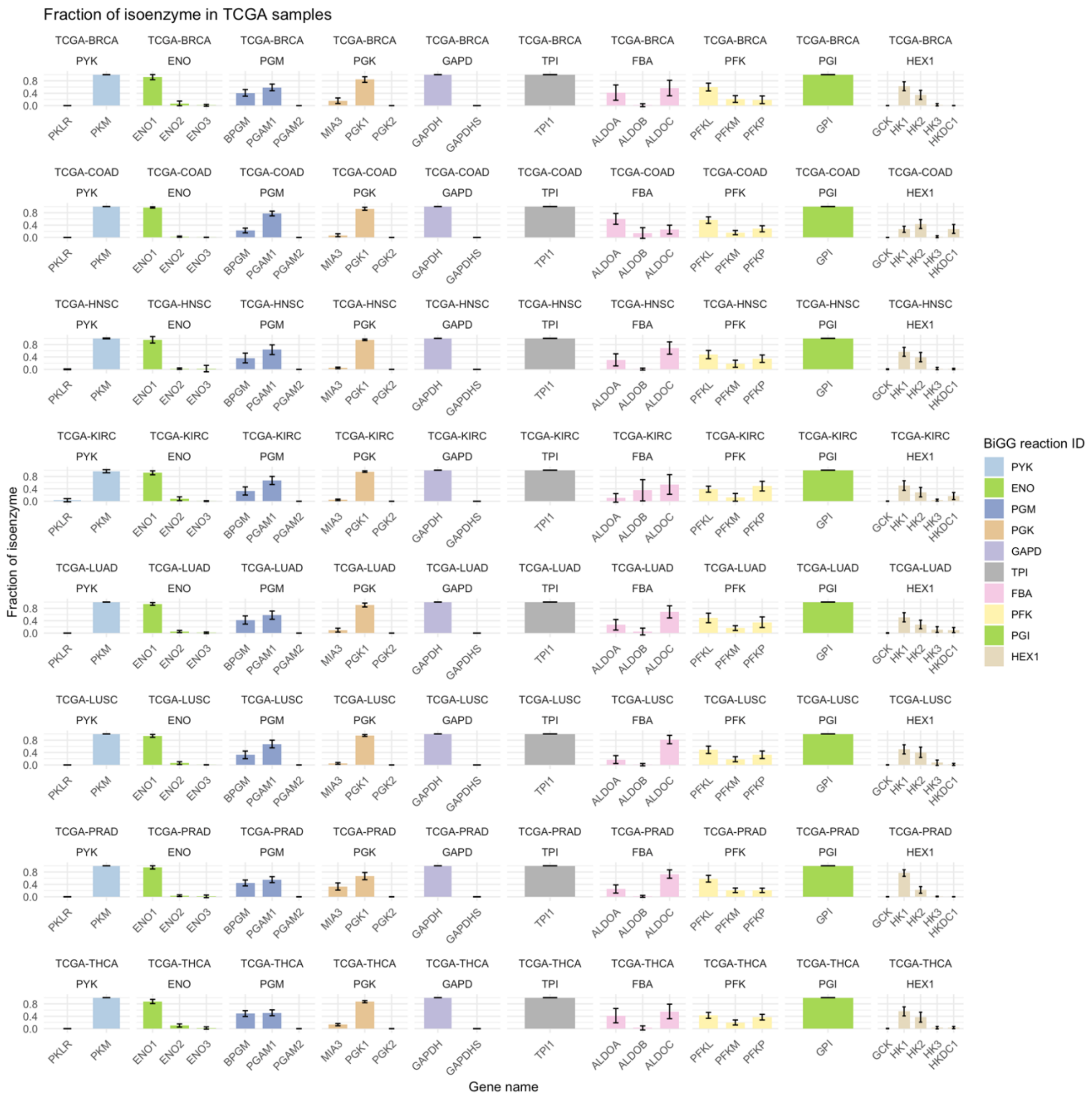


**Figure S4 (Related to Figure 3). Relative isozyme expression of glycolytic enzymes in the TCGA transcriptomics data.**

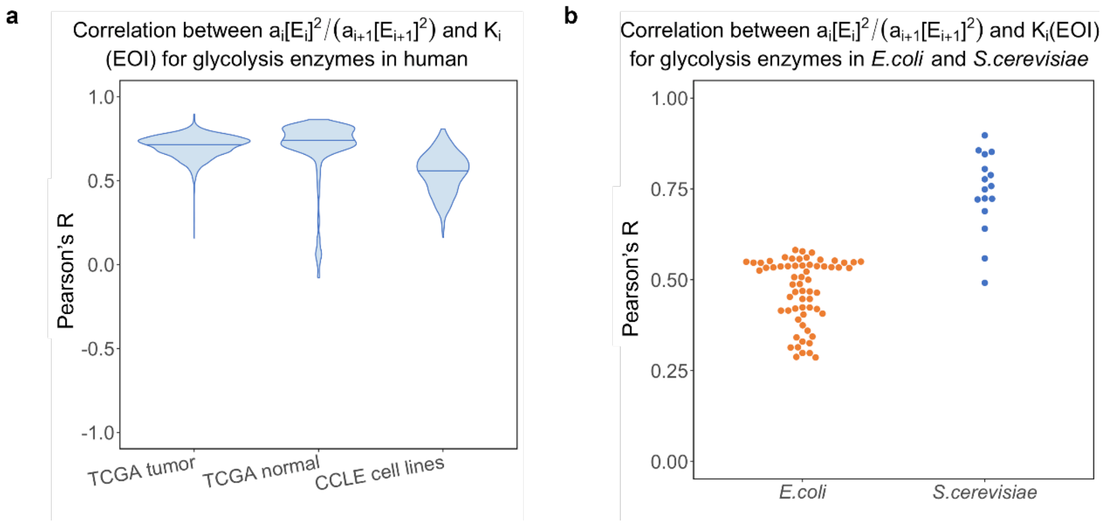


**Figure S5 (Related to Figure 4). Pearson correlation between K and CAQ for glycolytic enzymes.** (a) Distributions of the Pearson correlation between K and CAQ in the human datasets. (b) Distributions of the Pearson correlation between K and CAQ in the *E.coli* and *S.cerevisiae* datasets.

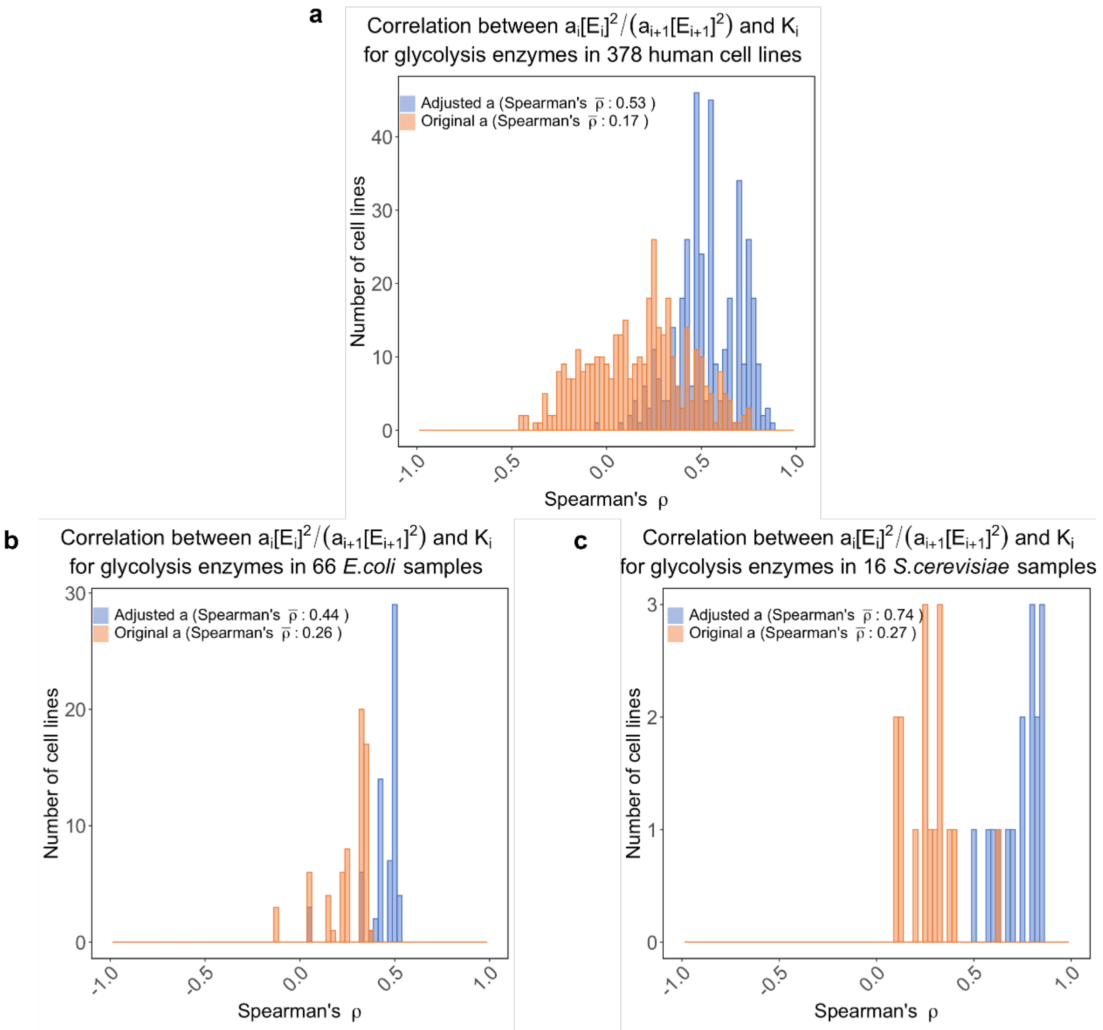


**Figure S6 (Related to Figure 4). Absolute metabolite concentrations are necessary for accurate estimation of catalytic efficiency and successful validation of the theory.** (a) Bar plot comparing the distributions of EOI computed for the human datasets based on catalytic efficiencies estimated with different methods. Original a: estimation of the catalytic efficiency $a$ does not incorporate substrate concentrations: $a = k_{cat}/K_m$; adjusted a: estimation of $a$ incorporates substrate concentrations: $a = \frac{k_{cat}}{K_m+[S]}$. (b) Same as in (a) but for the *E.coli* datasets. (c) Same as in (a) but for the *S.cerevisiae* datasets.

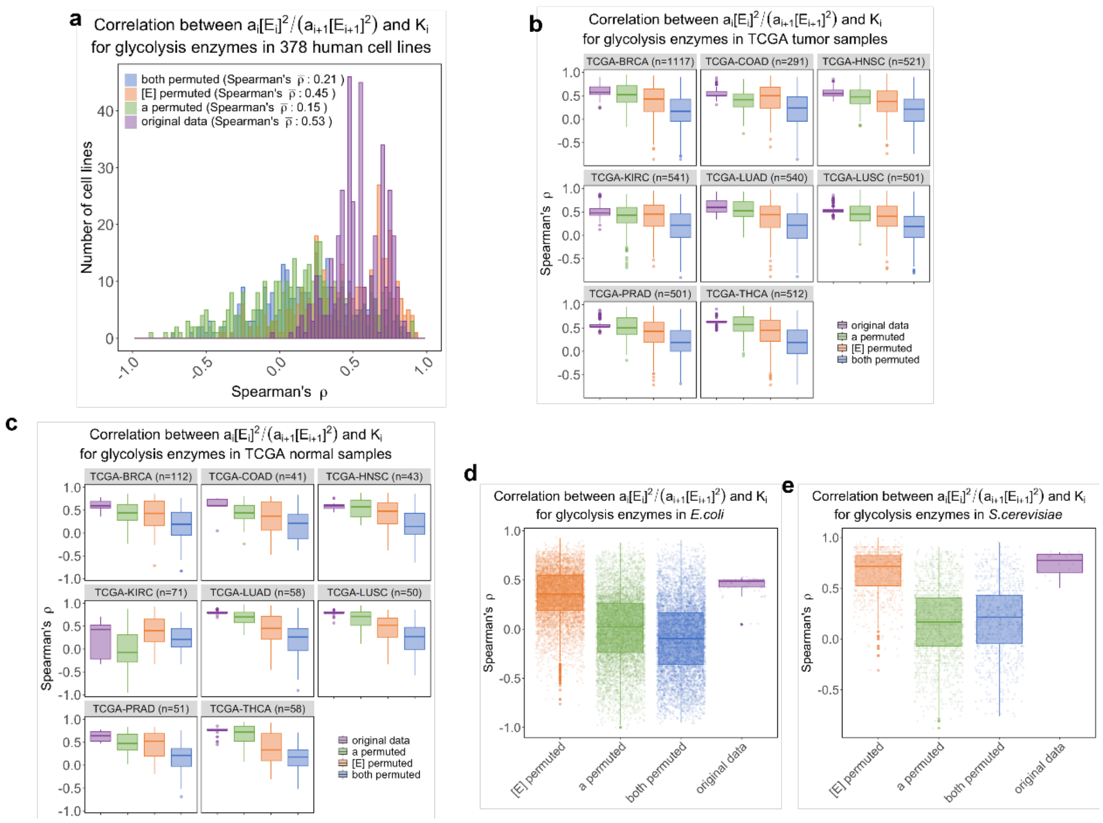


**Figure S7 (Related to Figure 4). Permutation tests on catalytic efficiency $a$ and enzyme abundance $[E]$.** (a) Bar graph comparing distributions of original EOI and those after individual or combinational permutation of catalytic efficiency $a$ and enzyme abundance $[E]$ for the human CCLE cell lines. (b) Box plots comparing distributions of original EOI and those after individual or combinational permutation of catalytic efficiency $a$ and enzyme abundance $[E]$ for the human TCGA tumor samples. (c) Same as in (b) but for human normal samples. (d) Same as in (b) but for the *E.coli* data. (e) Same as in (b) but for the *S.cerevisiae* data.

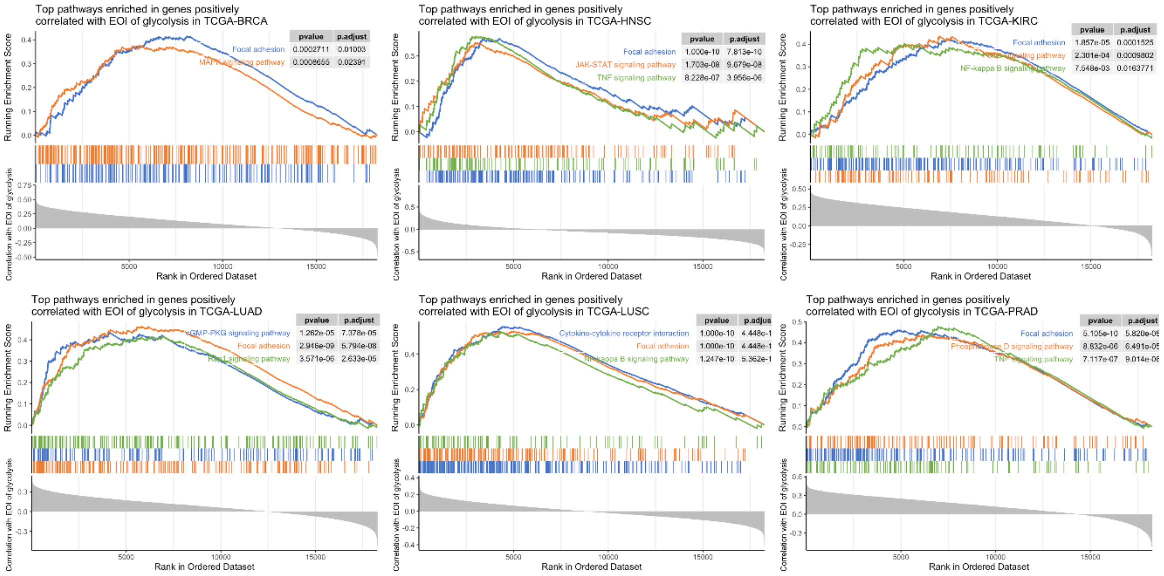


**Figure S8 (Related to Figure 5). GSEA plots showing top pathways enriched in genes positively correlated with EOI of glycolysis in different types of TCGA tumors.**

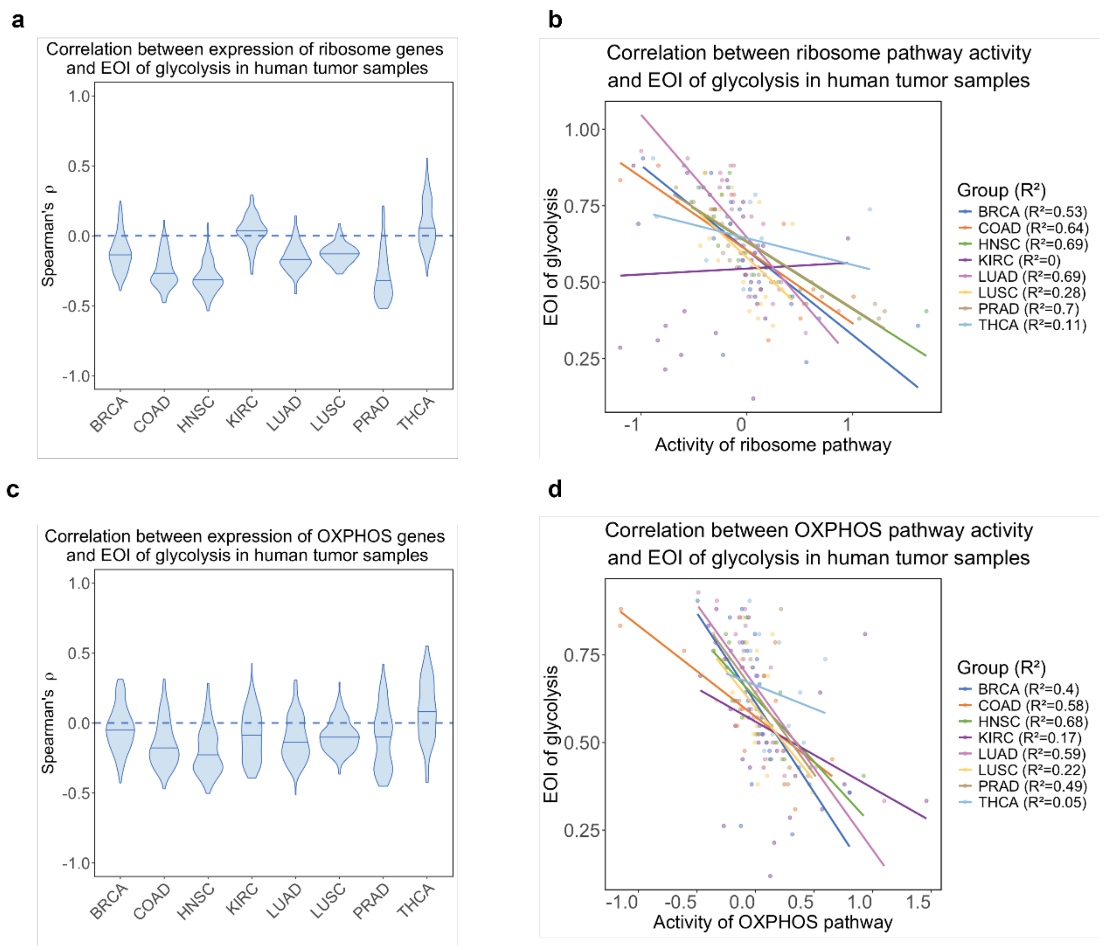


**Figure S9 (Related to Figure 5). Association of glycolytic EOI with expression of ribosome and OXPHOS genes.** (a) Violin plots showing distributions of Spearman's correlation coefficients between EOI of glycolysis and expression of ribosome genes. (b) Scatter plots correlating the EOI of glycolysis to activity of ribosome pathway defined as mean of Z-score normalized expression levels of ribosome genes. (c) Same as in (a) but for OXPHOS genes. (d) Same as in (b) but for OXPHOS genes.

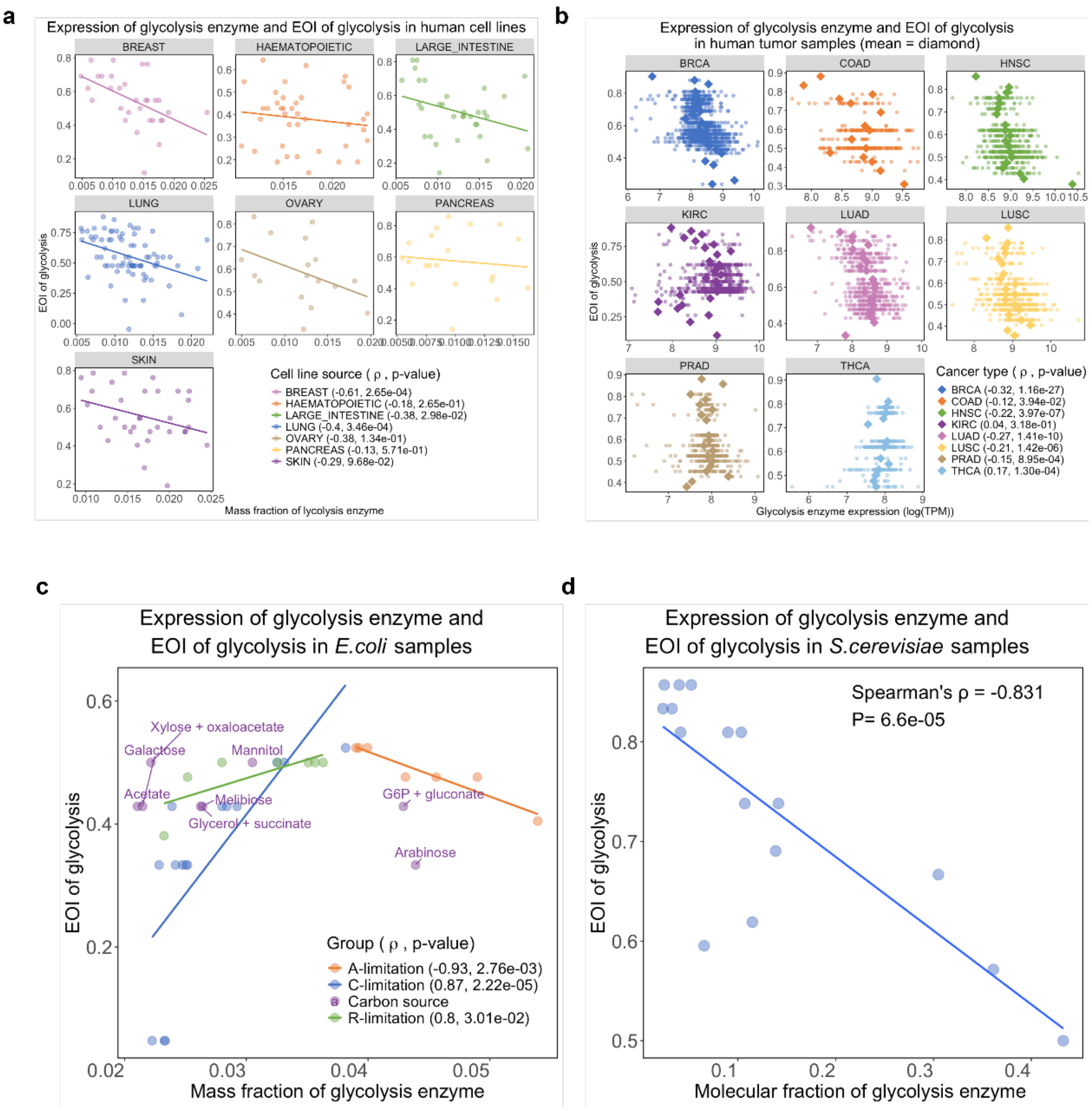


**Figure S10 (Related to Figure 5). Association between proteome investment of glycolysis and EOI of glycolysis.** (a) Scatter plots comparing mass fraction of glycolysis enzyme and EOI of glycolysis for human cell lines with different tissues-of-origin. (b) Same as in (a) but for human tumors in TCGA. (c) Same as in (a) but for *E.coli* samples. (d) Same as in (a) but for *S.cerevisiae* samples.

**Dataset S1**. Physiological intracellular conditions for *H.sapiens*, *S.cerevisiae*, and *E.coli*.

**Dataset S2**. Thermodynamic parameters for glycolytic reactions in *H.sapiens*, *S.cerevisiae*, and *E.coli*.

**Dataset S3**. Absolute concentrations of glycolytic intermediate metabolites in *H.sapiens*, *S.cerevisiae*, and *E.coli*.

**Dataset S4**. Kinetic parameters for glycolytic reactions in *H.sapiens*, *S.cerevisiae*, and *E.coli*.

### SI References

[1] Z. Dai and J. W. Locasale. Thermodynamic constraints on the regulation of metabolic fluxes. J Biol Chem, 293(51):19725–19739, 2018.

[2] Reinhart Heinrich and Edda Klipp. Control Analysis of Unbranched Enzymatic Chains in States of Maximal Activity. Journal of Theoretical Biology, 182(3):243–252, October 1996.
